# The fluid genomic organisation of jingmenviruses

**DOI:** 10.1101/2025.02.07.637040

**Authors:** Coralie Valle, Rhys H. Parry, Bruno Coutard, Agathe M.G. Colmant

## Abstract

Jingmenviruses are a distinct group of flavi-like viruses characterized by a genome consisting of four to five segments. Here, we report the discovery of three novel putative jingmenviruses, identified by mining publicly available metagenomics data from mosquito and arachnid samples. Strikingly, these novel jingmenvirus sequences contain up to six genomic segments, with pairs of homologous segments coding for putative structural proteins. Following this discovery, we found an additional homologous segment for two other jingmenvirus genomes, which had gone unnoticed in the initial publications. The presence of a single version of the segments coding for non-structural proteins suggests we have indeed identified jingmenviruses with infectious units that contain up to six segments. We compared these novel jingmenvirus sequences to published sequences, in particular the segments with multiple open reading frames, and we propose that the putative translation initiation mechanisms involved for these segments are ribosomal frameshift resulting in the fusion of open reading frames and leaky scanning for overlapping ORFs. These putative mechanisms, conserved for all jingmenvirus sequences analysed, including in homologous segments, require biological confirmation. We also generated structural models of two putative structural proteins in the duplicated segments, and the corresponding alignments enabled us to confirm or identify the homologous relationship between sequences that shared limited nucleotide or amino acid identity. Altogether, these results highlight the fluid nature of jingmenviruses, which is a hallmark of multipartite viruses. Different combinations of segments packaged in different virus particles could facilitate the acquisition or loss of genomic segments and a segment duplication following genomic drift. Our data therefore contribute to the evidence of the multipartite nature of jingmenviruses and the evolutionary role this organisation may play.

## Introduction

Jingmenviruses are related to flaviviruses but remarkably have a segmented genome, contrary to their close relatives (1). The first jingmenvirus identified was Jingmen tick virus (JMTV), from *Rhipicephalus microplus* ticks collected in China in 2010 (2). Since this initial discovery, dozens of other jingmenvirus genomes have been detected worldwide in a wide range of sample types (1). These sequences phylogenetically group into two main clades: one clade with sequences that are mostly tick- and vertebrate-associated and one clade with mainly insect-associated sequences (1). The best characterised insect-associated jingmenvirus species is Guaico Culex virus (GCXV), isolated from *Culex* mosquitoes collected in South and Central America between 2008 and 2012 and from *Culex* mosquitoes collected from Brazil in 2010 (3,4). GCXV has been found to have a replication restricted to insects both *in vitro* and *in vivo*, and there is evidence that this virus may have a multipartite organisation, meaning that viral particles encapsidate less fragments than the total number of genomic segments and multiple particles are required to initiate the replication cycle (3). Multipartite viruses are very common in plants and fungi, extremely rare in animals and have never been identified in bacteria (5). They seem to be more tolerant than their monopartite relatives to fluidity in their genomic organisation and to evolutive processes that facilitate functional gain by duplication or recombination (5).

The multi-segmented genomes of jingmenviruses are generally considered to include 4 segments except for GCXV, which has been found to have a fifth segment in most isolates but not all (in 4/6 sequenced isolates in Ladner *et al.*) (3). This fifth segment, named GCXV segment 5 encodes a viral protein called VP7, but no sequence homolog identified has been yet. Segment 5 is not essential for a productive viral infection and does not seem to be providing any fitness advantage *in vitro* or *in vivo* (3,6).

For most jingmenviruses, segment 1 encodes the non-structural protein NSP1 with RNA-dependant RNA polymerase (RdRp) and methyltransferase functional domains, similar to the non-structural protein NS5 of their close relatives in the *Orthoflavivirus* genus (7,8). Segment 2 encodes the putative glycoprotein VP1 or (VP1a and VP1b for some tick-associated jingmenviruses), and has a second open reading frame (ORF) corresponding to the putative small protein nuORF of unknown function(s) for sequences in the tick-associated jingmenvirus clade, or the putative structural protein VP4 for sequences in the insect-associated jingmenvirus clade. The glycoprotein VP1 in tick-associated sequences has recently been described as structurally homologous to the flavivirus envelope protein E, with no identified fusion loop homolog (9). Segment 3 encodes NSP2, with serine protease and helicase domains, similar to the flavivirus NS3 protein (10,11). Finally, segment 4 encodes putative structural proteins VP2 and VP3, with unknown functions.

This nomenclature is different for only two viruses, GCXV and its close relative Mole Culex virus (MoCV) isolated from *Culex* mosquitoes collected from Ghana in 2016 (12). For these two viruses, segment 1 encodes for NSP1, segment 2 for NSP2, segment 3 for putative structural proteins VP1, VP2 and VP3, segment 4 for putative structural proteins VP4, VP5 and VP6 and segment 5 for VP7 when present (1).

The segments of jingmenvirus genomes are capped at their 5’ end and polyadenylated at their 3’ end in the case of sequences in the tick-associated clade, making these fragments translation units, in which one to three coding sequences have been identified. The strategies used by jingmenviruses to ensure the translation of each coding sequence in the translation units, such as programmed ribosomal frameshifting or alternative translation initiation, have not been studied in detail to date (13).

Slippery heptanucleotides have been identified in a few jingmenvirus sequences, which suggests some ORFs are translated by a -1 ribosomal frameshift, the most common programmed frameshifting (GCXV and MoCV between ORFs coding for VP1 and VP3 as well as VP5 and VP6; in JMTV, Alongshan virus and Wuhan cricket virus between ORFs coding for VP2 and VP3; in ALSV between ORFs coding for VP1a and VP1b) (3,12,14,15). These predictions are yet to be confirmed biologically, with the exception of GCXV frameshift between VP1 and VP3, for which the -1 ribosomal frameshift products have been confirmed by mass spectrometry. Other than these instances of likely ribosomal frameshift, no other mechanism has been suggested for the translation of multiple ORFs in jingmenvirus genomic segments, despite the existence of overlapping ORFs, possibly translated through a cap-dependent deviant translation initiation, a mechanism commonly described in viruses allowing the production of multiple functions from a single RNA (13).

To date, it is unclear how and if the segmented and putative multipartite nature of jingmenviruses confers them a selective advantage over their monopartite relatives. In order to gain knowledge on these viruses, we have searched for novel jingmenvirus sequences using publicly available next-generation sequencing data. We have also analysed the sequence of the genomic segments of 25 jingmenvirus species to better understand their organisation. The data generated lead us to propose multiple new possible genomic organisations for jingmenviruses, with variable numbers of segments.

## Material and methods

### Discovery of new mosquito-related jingmenvirus sequences

We used the online Serratus platform, specifically the RdRp search functionality, to find novel mosquito-derived jingmenvirus sequences in published RNA sequencing datasets (16). We searched the “Family” entitled Unclassified-899, as we identified that this corresponded to insect-related jingmenviruses in Serratus. The filters used were a percentage identity above 75% and a score above 50. For the 97 matches found, we selected the ones with a mosquito or mosquito-related host, as we were originally looking for novel mosquito-derived jingmenvirus sequences in this study.

Additionally, we screened for divergent hits related to the Flavi-like segmented virus strain US001 NS5 gene (MN811583.1) and manually assembled new sequences from SRA datasets with scores above 15, percentage identity above 50% and more than 25 reads (17).

We either downloaded the raw sequencing reads from the NCBI website and assembled the new genomic sequences using Geneious Prime 2024.0.7 or MEGAHIT v1.2.9; or we performed the assemblies using NCBI BLAST with its direct access to the Sequence Read Archive (SRA) database, with the Megablast algorithm optimised for highly similar sequences with maximum target sequences set to the highest setting (5000). In both cases, we used the RdRp contigs assembled by Serratus or published sequences (GCXV KM461666 to KM461670; SAIV7 KR902717 to KR902720; OKIAV332 MW314687) as references for the assemblies.

### Sequence analysis

Multiple sequence alignments were performed with MUSCLE using Geneious Prime 2024.0.7, which provided the percentage identity matrices.

The best models to use in order to infer the maximum likelihood phylogenies presented here were selected for each alignment using the online tool Smart Model Selection in PhyML (http://www.atgc-montpellier.fr/sms/ (18)). The phylogenetic analyses were performed using the PhyML plugin in Geneious Prime 2024.0.7, with an LG substitution model and 100 bootstraps. Phylogenies were mid-point rooted using FigTree (http://tree.bio.ed.ac.uk/software/figtree/).

### Identification of slippery sequences and ribosomal frameshifts

We followed guidelines described by McNair *et al.* to identify potential heptanucleotide slippery motifs that could result in a ribosomal frameshift in jingmenvirus segments with multiple ORFs (19). We used HotKnots to compute potential secondary structures in the 100 nucleotides following the identified slippery heptanucleotides and to estimate the minimum free energy of said secondary structures (www.rnasoft.ca) (20). We then used the PRFECT software developed by McNair *et al.* and confronted the results obtained to the motifs identified above. When multiple possible motifs were identified, we kept the ones identified using both the manual method and the software results. When multiple motifs remained, we selected the one with the highest probability value generated by the PRFECT software. When this criterion did not help to discriminate between multiple motifs, or when PRFECT had not identified a motif in the sequence, although we had identified several, we performed amino acid sequence alignments with the multiple resulting protein sequences. We selected the frameshift position that resulted in the most conserved amino acid sequence. All predicted amino acid sequences were then translated and aligned to confirm that the resulting frameshifted sequences shared a similar sequence homology with related sequences as the rest of the protein, and that highly conserved residues were translated correctly.

### Screening for IRES sequences in jingmenvirus genomic segments with multiple ORFs

We used the tool DeepIRES developed by Zhao *et al.* to screen the segments with multiple ORFs of jingmenvirus genomes for the presence of internal ribosome entry site (IRES) (21). We then compared the prediction obtained from DeepIRES with the predicted ORFs in these genomic segments to see if the DeepIRES predictions could be associated to a biological relevance.

### Protein structure modelling

The structures of VP4 and VP1 of GCXV, and the mosquito-derived jingmenvirus sequences discovered here; as well as both VP2 versions of SAIV7 were predicted using AlphaFold2 Google Colab (22). Model super-impositions were generated using Flexible structure Alignment by Chaining Aligned fragment pairs, allowing Twists (FATCAT) server (23) and visualized using UCSF Chimera X (24). The similarity detected by FATCAT is evaluated by a p-value. The smaller the p-value, the more statistically significant the similarity between the two structures. Pairwise sequence alignments were performed using the Needleman-Wunsch algorithm provided by UCSF Chimera X and analysed with ESPript (25).

## Results

### Discovery of new mosquito-related jingmenvirus sequences

Using Serratus to search for novel mosquito-derived jingmenvirus RdRp coding sequences, we identified seven datasets of interest. Five datasets (SRX8323447, SRX8323446, SRX8323444, SRX8323437, SRX8323431) from wild-caught mosquito excreta in Far North Queensland, Australia, in 2018 had a partial match with 76-78% nucleotide identity to the RdRp of GCXV and MoCV (26). One dataset (SRX4113301) from a pool of 4,400 *Cx. tritaeniorhynchus* mosquitoes collected from Yunnan, China, in 2016 had a partial match with 79-80% nucleotide identity to the RdRp of GCXV and MoCV (27). The seventh dataset (SRX833670) from an uncultured mosquito pool collected from Zhejiang, China, in 2013 had a match with 99% identity to the RdRp of Shuangao insect virus 7 (SAIV7). This mosquito pool contained the following species: *Aedes albopictus, Armigeres subalbatus, Anopheles paraliae, An. sinensis, Culex pipiens, Cx. sp., Cx. tritaeniorhynchus* (28).

We first focused on the most divergent sequences and assembled the novel jingmenvirus genomic sequences from mosquito excreta and *Cx. Tritaeniorhynchus*, then moved on to the SAIV7 positive dataset. All newly described sequences were deposited in Genbank under accession numbers XXXXXXX-XXXXXXXX (submitted).

### New jingmenvirus sequences from mosquito excreta

The assemblies performed on the mosquito excreta datasets identified with Serratus yielded the genomic sequence for a putative novel jingmenvirus, most closely related to GCXV according to NCBI BLASTx search results. The genome was first assembled using SRX8323431 and all segments were subsequently detected in the other four datasets identified with Serratus. Strikingly, we identified six putative viral segments instead of five, as is the case for GCXV or four, as is the case for all other described jingmenviruses.

The full nucleotide sequences of segments 1 (NSP1) and 2 (NSP2) shared only 71% and 50% identity, respectively, with GCXV segments 1 and 2. Among the other four segments, two putative versions of both segments 3 (34.9% and 50.4% nucleotide identity with GCXV) and 4 (49.7% and 48.2% nucleotide identity with GCXV) were assembled from the data (Figure 1), while no sequence related to GCXV segment 5 was identified. We searched for evidence of co-infection or multiple infections in the datasets by searching for additional segments 1 and 2. We did not find any in the data, even when assemblies were performed with less stringent mapping criteria, using GCXV and the new sequences as references, suggesting the six segments participate in a single infection unit.

**Figure 1:**
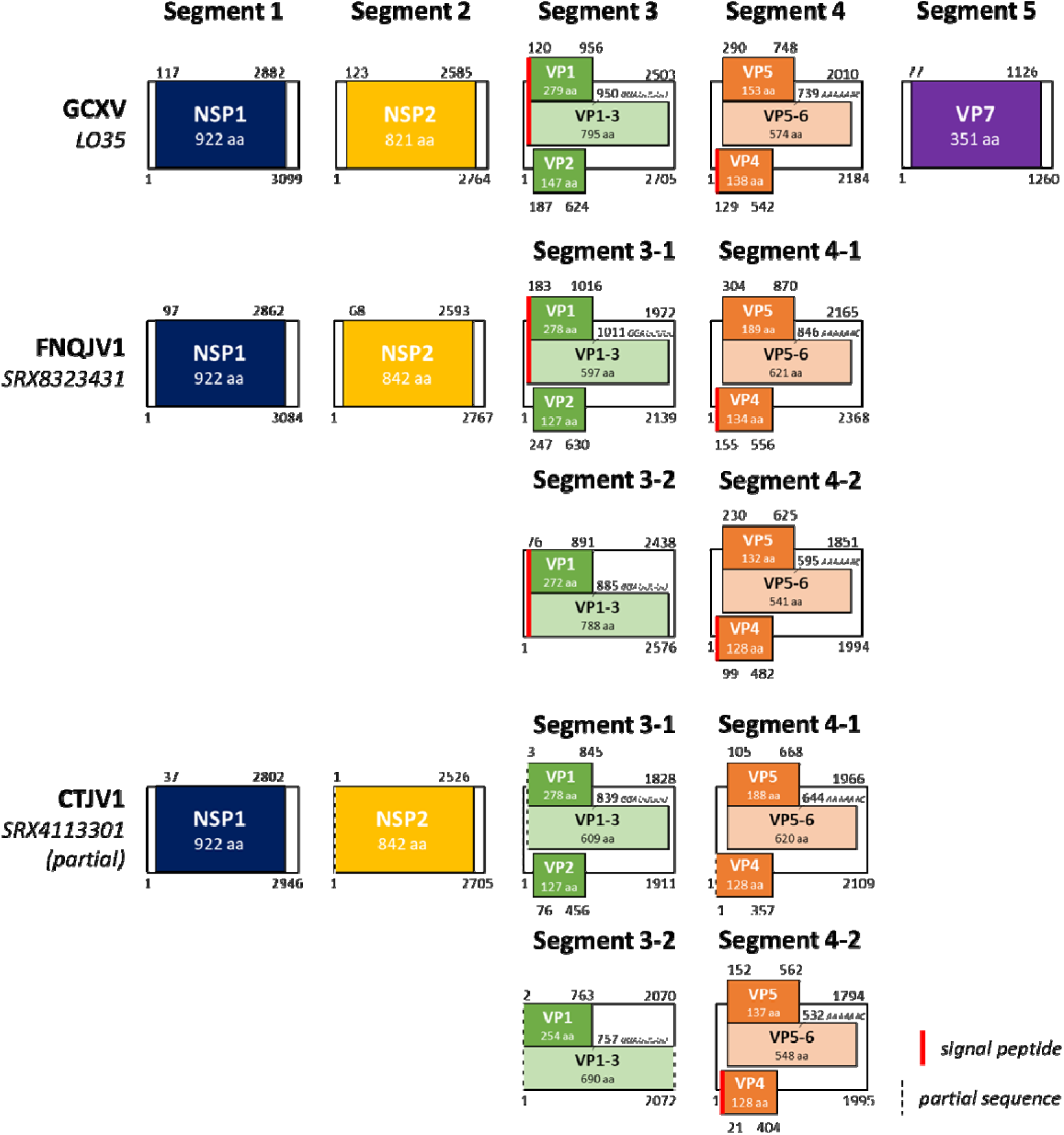
Genomic organisation of two new mosquito-derived jingmenviruses, FNQJV1 and CTJV1 compared to the published sequence of GCXV. ORFs are represented by blocks of colour, and translation resulting from a ribosomal frameshift is represented by blocks in pastel colours, with the position of the frameshift and slippery heptanucleotide sequence specified underneath.

In the two segments designated segment 3, three ORFs coding for VP1 to VP3 were identified in the first one (hereafter designated segment 3-1), similarly to GCXV. Only two ORFs, coding for VP1 and VP3 were identified in the second one (hereafter designated segment 3-2).

In segment 3-1, most of the genetic distance was attributed to the second half of the sequence (Figure 2). Indeed, VP1 and VP2 coding sequences were identified (50% nucleotide identity with GCXV) whereas the VP3 coding sequence only shared 24% nucleotide identity with GCXV VP3, and was almost 600 nucleotides shorter than the GCXV VP3 ORF, while the 3’UTR of this segment (163 nucleotides) had a length similar to that of GCXV segment 3 3’UTR (199 nucleotides), indicating the segment is indeed shorter due to the VP3 ORF length.

**Figure 2:**
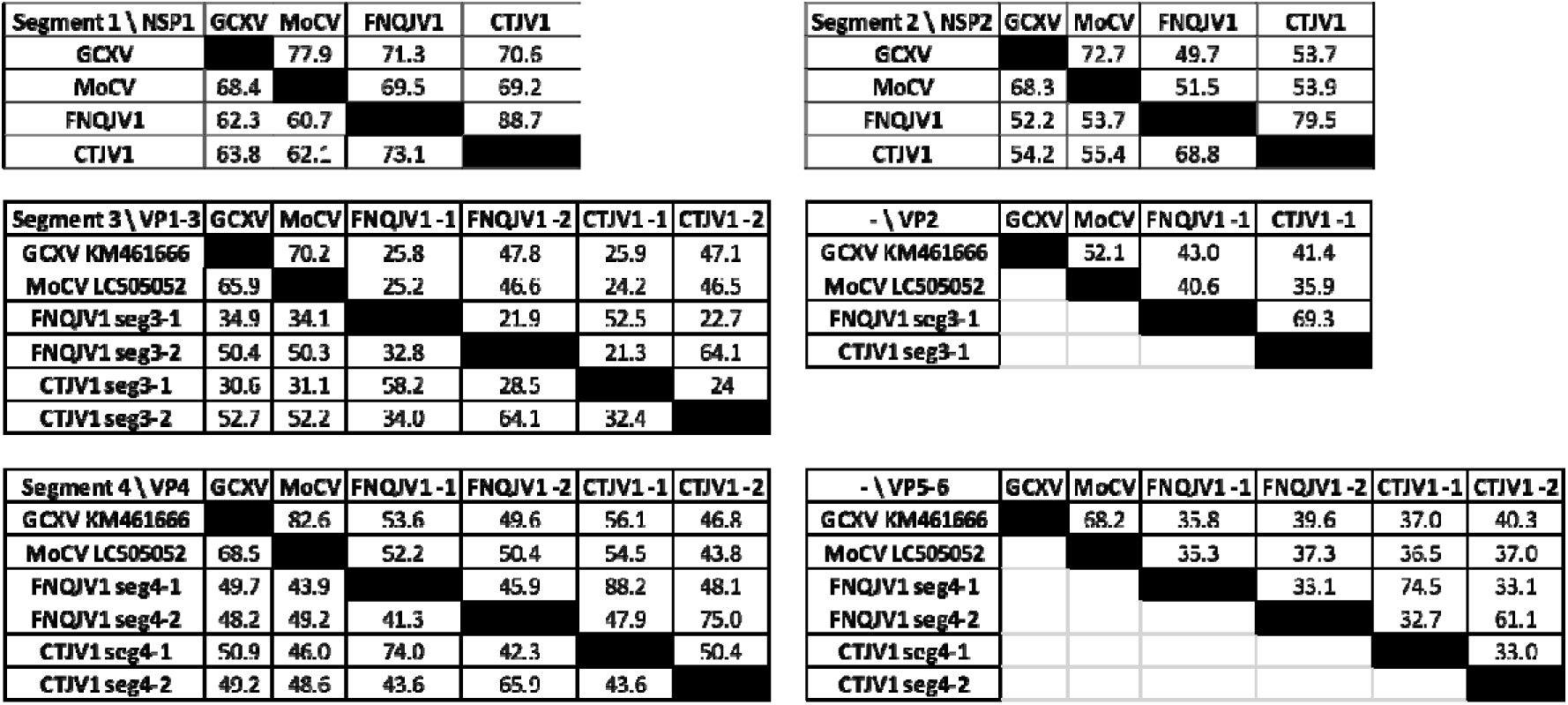
Percentage identity derived from multiple sequence alignment of GCXV, MoCV, FNQJV1 and CTJV1 genomic sequences. The bottom left of each table represents nucleotide identity over the whole genomic segment. The top right of each table corresponds to amino acid percentage identity over the corresponding ORF.

The other version of segment 3 (segment 3-2) shared 50% nucleotide identity overall with GCXV and 33% nucleotide identity with the first version in segment 3-1. While ORFs for VP1 and VP3 were identified, no obvious additional ORF with START and STOP codon or possible frameshifting with VP1 and VP3 was identified, thus VP2 coding sequence is not present in this version.

We chose the designations segment 3-1 and 3-2 based on the similarity of genomic organisation compared to GCXV, with segment 3-1 being the version with the genomic organisation closest to GCXV.

The two versions of segment 4 shared 41% nucleotide identity with each other and 48-50% nucleotide identity with GCXV. ORFs coding for VP4 and VP5-6 were detected in both versions. Considering that the genomic organisation was the same for GCXV and these two segments, we chose to assign numbers to the versions based on the percentage nucleotide identity with GCXV over the whole segment. Segment 4-1 is slightly more similar to GCXV than segment 4-2 with 49.7% nucleotide identity, *versus* 48.2% (Figure 2).

We searched the 52 SRA datasets from this BioProject (PRJNA631724) for each of the six identified segments using NCBI BLAST. We found highly similar reads mapping to these jingmenvirus-associated sequences in 11 datasets, all from Far North Queensland, none from South East Queensland. We tentatively named this group of six sequences Far North Queensland jingmenvirus 1 (FNQJV1). The six segments were always found together, except in two datasets where the number of identified reads were below 50 for the most detected segments, therefore, which might not have had sufficient levels of viral RNA for the whole genome to be detected (Table 1). Of note, segments 3-2 and 4-2 were present in higher numbers of reads than 3-1 and 4-1, respectively in the majority of datasets.

**Table 1:** Number of reads mapping to FNQJV1 six genomic segment sequences in datasets from BioProject PRJNA631724 and to CTJV1 in datasets from BioProject PRJNA472635. The assemblies were performed using BLAST NCBI with the megablast algorithm, optimised for highly similar sequences, with a maximum target sequence set to the highest setting (5000). This method did not detect reads similar to FNQJV1 sequences in any of the other 41 datasets from PRJNA631724, nor did it detect reads similar to CTJV1 in the other two datasets from PRJNA472635. In bold are the datasets that had been identified using Serratus, and underlined is the dataset used to assemble FNQJV1 and CTJV1 segment sequences. seg: Segment.

| Dataset | Reference | seg1 | seg2 | seg3-1 | seg3-2 | seg4-1 | seg4-2 |
| --- | --- | --- | --- | --- | --- | --- | --- |
| SRX8323428 | FNQJV1 | 156 | 162 | 219 | 358 | 119 | 380 |
| **<u>SRX8323431</u>** | FNQJV1 | >5000 | >5000 | >5000 | >5000 | >5000 | >5000 |
| SRX8323435 | FNQJV1 | 66 | 181 | 275 | 487 | 318 | 535 |
| SRX8323436 | FNQJV1 | 92 | 254 | 65 | 248 | 107 | 246 |
| **SRX8323437** | FNQJV1 | 698 | 1389 | 1727 | 2322 | 1311 | >5000 |
| SRX8323439 | FNQJV1 | 0 | 42 | 3 | 26 | 11 | 15 |
| SRX8323443 | FNQJV1 | 1 | 3 | 4 | 6 | 2 | 0 |
| **SRX8323444** | FNQJV1 | 775 | 1428 | 601 | 1655 | 610 | 2052 |
| **SRX8323446** | FNQJV1 | 991 | 1216 | 923 | 1951 | 1357 | 2532 |
| **SRX8323447** | FNQJV1 | 2566 | 3546 | 3450 | >5000 | 3836 | >5000 |
| SRX8323477 | FNQJV1 | 36 | 33 | 62 | 199 | 21 | 835 |
| **<u>SRX4113301</u>** | CTJV1 | 357 | 132 | 241 | 193 | 135 | 3196 |

### New jingmenvirus sequence from *Culex tritaeniorhynchus*

Another dataset of interest identified with Serratus was from a pool of *Cx. tritaeniorhynchus* mosquitoes collected from Yunnan, China in 2016 (SRX4113301). From this dataset, we obtained the genomic sequence for another novel jingmenvirus, Culex tritaeniorhynchus jingmenvirus 1 (CTJV1), and despite a lower coverage for the assemblies than for FNQJV1, we identified that CTJV1 and FNQJV1 share a common genomic organisation. Indeed, two versions of segments 3 and 4 were assembled from the data, closely related to the corresponding versions of FNQJV1 (one version of segment 3 has three putative ORFs for VP1 to VP3 and the second version has no VP2 ORF; both versions of segment 4 have three putative ORFs), while only one version of segments 1 and 2 was assembled for each (Figure 1, Figure 2).

Using NCBI BLAST, we searched the three SRA datasets from this BioProject (PRJNA472635) for each of the six identified CTJV1 segments and found reads mapping to CTJV1 only in the original *Cx. tritaeniorhynchus* dataset identified with Serratus (Table 1).

Interestingly, the two newly identified mosquito-derived sequences (FNQJV1 and CTJV1) are more closely related to each other than to any other published sequence, for all segments, despite the geographical distance of the samples that yielded the positive datasets (Figure 2). Indeed, FNQJV1 and CTJV1 share 89% amino acid identity over NSP1, 80% over NSP2, 53% over VP1/VP3 and 69% over VP2, for segment 3-1, 64% over VP1/VP3 for segment 3-2, 88% and 75% over VP4 for segment 4-1 and 4-2 respectively, 75% and 61% over VP5-6 for segment 4-1 and 4-2 respectively (Figure 2). These values are much higher than the percentage identity they shared with GCXV or MoCV, their closest published relative. Moreover, in addition to their shared genomic organisation, FNQJV1 segment 3-1 is much more closely related to its homolog CTJV1 segment 3-1 than it is to FNQJV1 segment 3-2. This relationship is true for all homologous segments and can be observed in the phylogenetic analyses presented in Figure 3. It should be noted that the lowest percentage identity values were observed for segment 3 and in particular for VP1-3. These low values are heavily influenced by the divergence of the VP3 sequence in segments 3-1.

**Figure 3:**
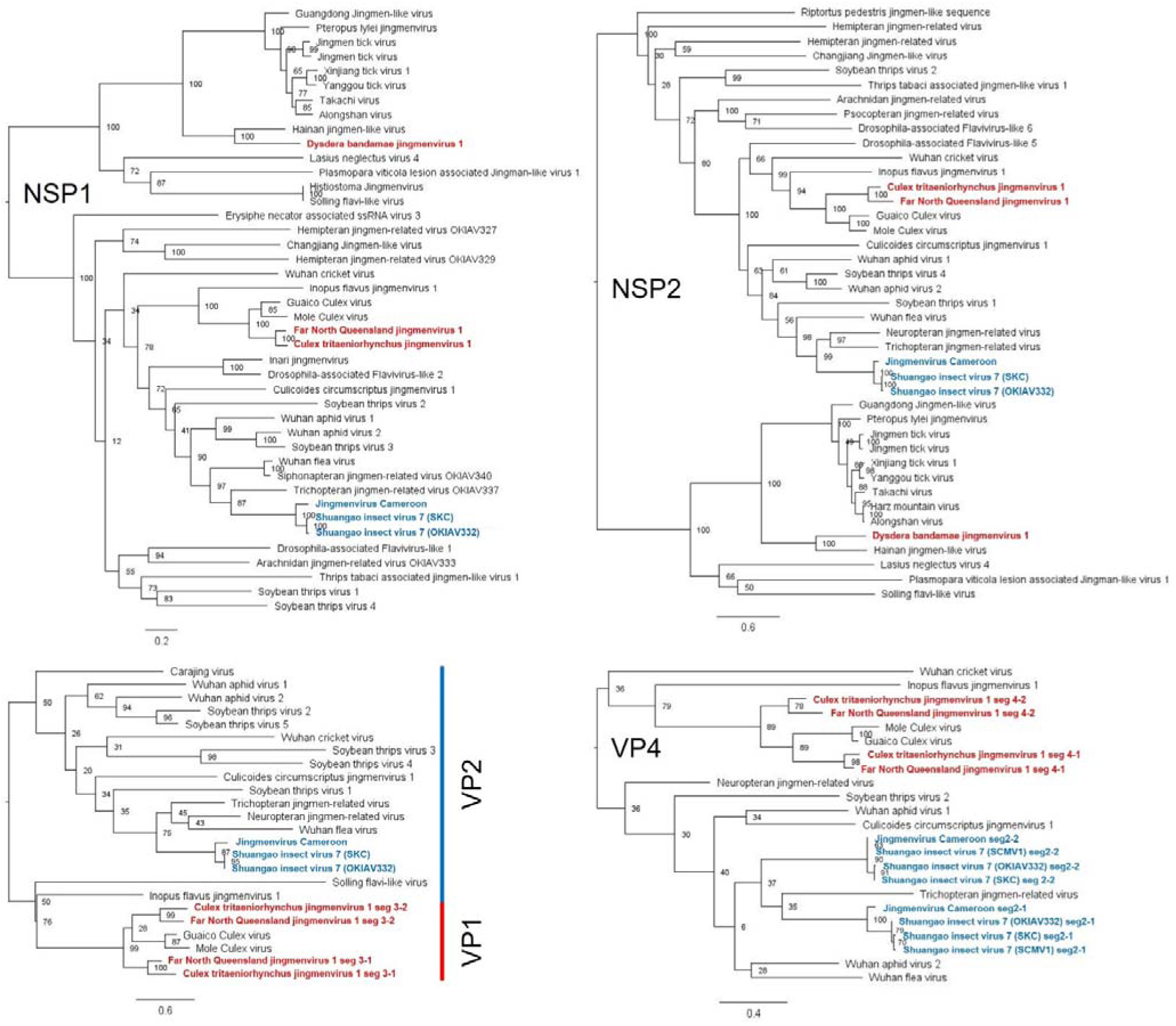
Maximum likelihood analysis of homologous jingmenviruses amino acid sequences: NSP1, NSP2, VP1 (mosquito-derived sequences) or VP2 (insect-derived sequences) and VP4 (mosquito- and insect-derived). The phylogenetic trees were constructed using the LG model, gamma distributed with 100 bootstraps (branch labels) and midpoint rooted using FigTree. The scale bar represents the number of amino acid substitution per site. The sequences highlighted in red were discovered and described in this study, the sequences highlighted in blue are part of viruses for which we identified additional segments.

### Identification of an additional genomic segment for Shuangao insect virus 7

The last dataset of interest identified with Serratus was an uncultured mosquito pool collected from Zhejiang, China, in 2013 (SRX833670). The assemblies performed on these data yielded the genomic sequence (segments 1, 2, 3, 4) for a new strain of SAIV7. These nucleotide sequences share 99.3 to 99.5% identity to the SAIV7 reference sequences (KR902717 to KR902720), obtained from another uncultured insect pool (SRX833685) from the same BioProject (PRJNA271540) (14).

When comparing these four sequences to published sequences using BLAST, we noticed that segments 1 (coding for NSP1), 3 (NSP2) and 4 (VP2 and VP3) shared 98% identity with Dipteran jingmen-related virus (OKIAV332) (MW314686, MW314688, MW314689), while segment 2 (coding for VP1 and VP4, thereafter designated segment 2-1) shared only 43% nucleotide identity with OKIAV332 segment 2 (MW314687; thereafter designated segment 2-2) (Figure 4) (29). Moreover, SAIV7 segment 2-1 shared 99% identity with Sichuan mosquito virus 1 (SCMV1) segment 2, this segment being the only published sequence for this virus (MZ556307) (30). In order to determine whether these datasets were proof of segment 2 reassortment between viruses or an indication that this virus genomic sequence actually includes five segments, we searched for the five sequences (segments 1, 2-1, 2-2, 3 and 4) using NCBI BLAST/SRA in these records as well as in the dataset identified with Serratus (SAIV7: SRX833685; OKIAV332: SRX798056; SCMV1: SRX10979868; Serratus: SRX833670). We detected all five segments in the four datasets (Table 2), with VP1 and VP4 ORFs in both segment 2-1 and 2-2 (Figure 4).

**Figure 4:**
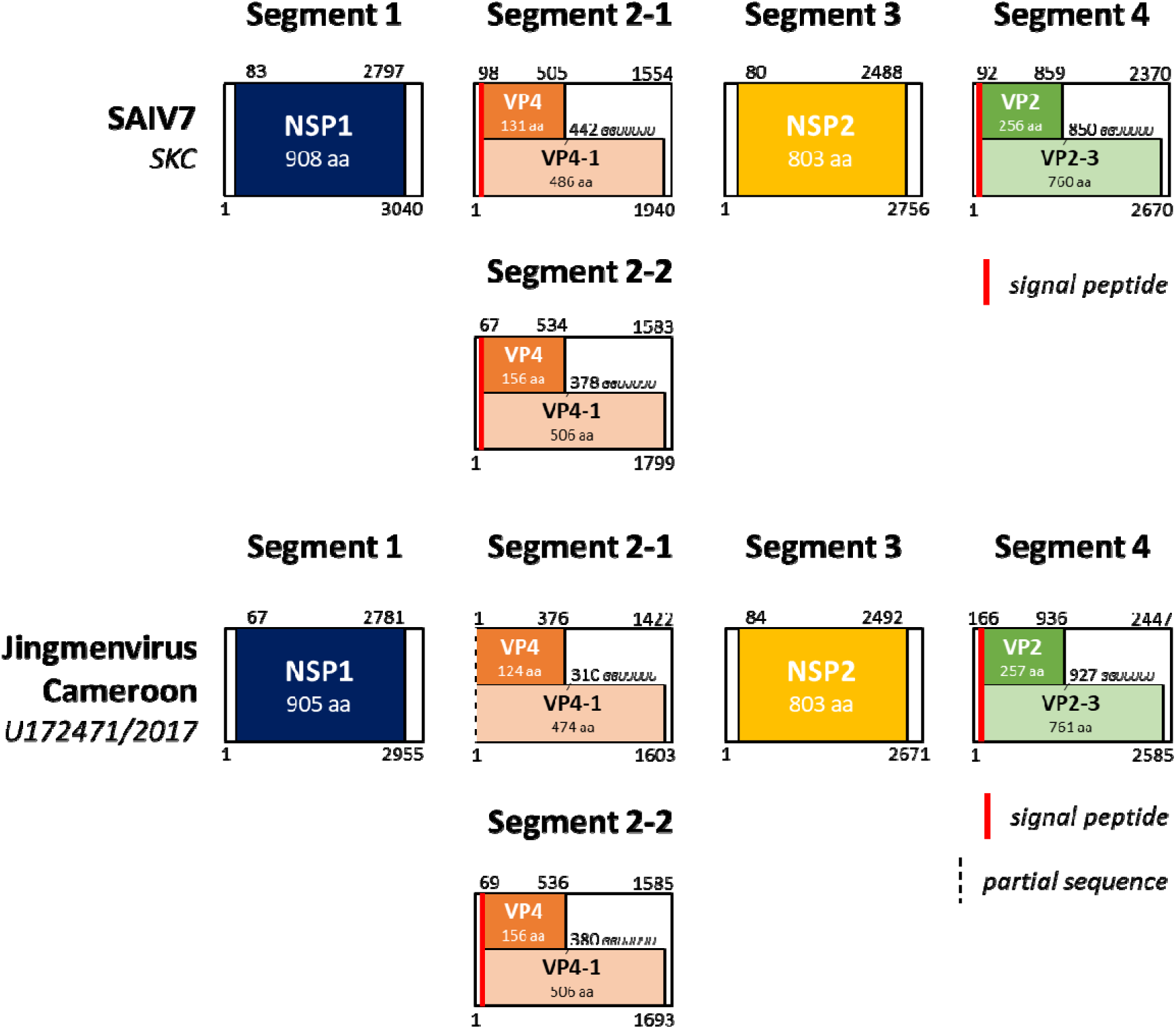
Genomic organisation of Shuangao insect virus 7 (SAIV7) and Jingmenvirus Cameroon (JVC). ORFs are represented by blocks of colour, and translation resulting from a ribosomal frameshift is represented by blocks in pastel colours, with the position of the frameshift and slippery heptanucleotide sequence specified underneath.

**Table 2:** Number of reads mapping to the five genomic segment sequences of SAIV7 in the datasets of the three studies with published sequences highly similar to SAIV7, and in the datasets identified with Serratus. The assemblies were performed using BLAST NCBI with the megablast algorithm, optimised for highly similar sequences, with a maximum target sequence set to the highest setting (5000)nn. The references used for the BLAST search were KR902717 (segment 1), KR902718 (segment 2-1), MW314687 (segment 2-2), KR902719 (segment 3) and KR902720 (segment 4). seg: Segment.

| Dataset | seg1 | seg2-1 | seg2-2 | seg3 | seg4 | Sample | Country | BioProject |
| --- | --- | --- | --- | --- | --- | --- | --- | --- |
| SRX833685 (SAIV7) | >5000 | >5000 | >5000 | >5000 | >5000 | *Chrysopidae sp., Psychoda alternata, Diptera sp.* | China | PRJNA271540 |
| SRX798056 (OKIAV332) | 859 | 888 | 862 | 682 | 673 | *Clogmia albipunctata* | USA | PRJNA267928 |
| SRX10979868 (SCMV1) | 20 | 96 | 7 | 60 | 28 | Mosquito | China | PRJNA680461 |
| SRX833670 (Serratus) | 553 | 2959 | 426 | 891 | 562 | Mosquito | China | PRJNA271540 |
| SRX5887211 | 713 | 1325 | 736 | 837 | 1036 | *Anser indicus* | India | PRJNA526291 |
| SRX13680292 | 991 | >5000 | 1547 | 1693 | 1507 | *Hystrix brachyura* | China | PRJNA795267 |
| SRX13667241 | >5000 | >5000 | >5000 | >5000 | >5000 | *Hystrix brachyura* | China | PRJNA795267 |
| SRX13680277 | 1135 | >5000 | 4253 | 2355 | 3864 | *Paguma larvata* | China | PRJNA793740 |
| SRX14333807 | 2506 | 4915 | 1137 | 4092 | >5000 | Wastewater | USA | PRJNA771693 |
| SRX14333808 | 2321 | 2625 | 732 | 2760 | 4744 | Wastewater | USA | PRJNA771693 |
| SRX14333809 | 1544 | 3061 | 502 | 2147 | 2627 | Wastewater | USA | PRJNA771693 |
| SRX3035916 | 357 | 2757 | 569 | 684 | 580 | Wastewater | Brazil | PRJNA395784 |

Furthermore, using Serratus looking specifically for SAIV7-related sequences, we found 12 datasets that had RdRp sequences with >95% identity to SAIV7 (SRX5887211, SRX13680292, SRX13667241, SRX833670, SRX833685, SRX13680277, SRX14333774, SRX14333807, SRX14333808, SRX14333809, SRX16702048, SRX3035916). We had already screened two of these datasets (SRX833670 and SRX833685) for the presence of the five segments. Two other datasets (SRX14333774 and SRX16702048) were not available for assembly with NCBI BLAST/SRA and were not included in the further search. We found that the five genomic segments (1, 2-1, 2-2, 3 and 4) were represented in the remaining 8 datasets. The positive datasets originated from samples as diverse as an oral swab of bar-headed goose *Anser indicus* from India, a nasal swab and faecal sample from Malayan porcupine *Hystrix brachyura* from China, a faecal sample from masked palm civet *Paguma larvata* from China and wastewater from the USA and Brazil (Table 2).

In this study, we therefore found five SAIV7 genomic segment sequences in a wide distribution of hosts: a sample with pooled *Chrysopidae sp., Psychoda alternata, Diptera sp.;* a sample with pooled *Ae. albopictus, Ar. subalbatus, An. paraliae, An. sinensis, Cx. pipiens, Cx. sp, Cx. tritaeniorhynchus;* and samples containing each *Clogmia albipunctata, Hystrix brachyura* swabs*, Paguma larvata* swabs*, Anser indicus* swabs and waste water; in samples collected from China, India, Brazil and the USA between 2012 and 2021 (28–31).

### Identification of an additional genomic segment for Jingmenvirus Cameroon

As we were comparing the SAIV7 genomic sequences to published sequences using NCBI BLAST, we noticed that their most closely related sequences are the genomic segments of a jingmenvirus sequence detected from human blood samples from Cameroon (70-83% nucleotide identity for segments 1, 2-1, 3 and 4) (Figure 5) (32). The authors reported 4 genomic segments for this virus they named Jingmenvirus sp. strain Cameroon/U172471/2017 (hereafter described as Jingmenvirus Cameroon or JVC). Upon our request, the raw sequencing reads for this metagenomics study were added to the NCBI SRA public database (SRR31402545, SRR31402546, SRR31402547) and we were able to assemble a fifth genomic segment from the raw data, highly similar to SAIV7 segment 2-2 (93-99% nucleotide identity to SAIV7 segment 2-2) (Figure 5).

**Figure 5:**
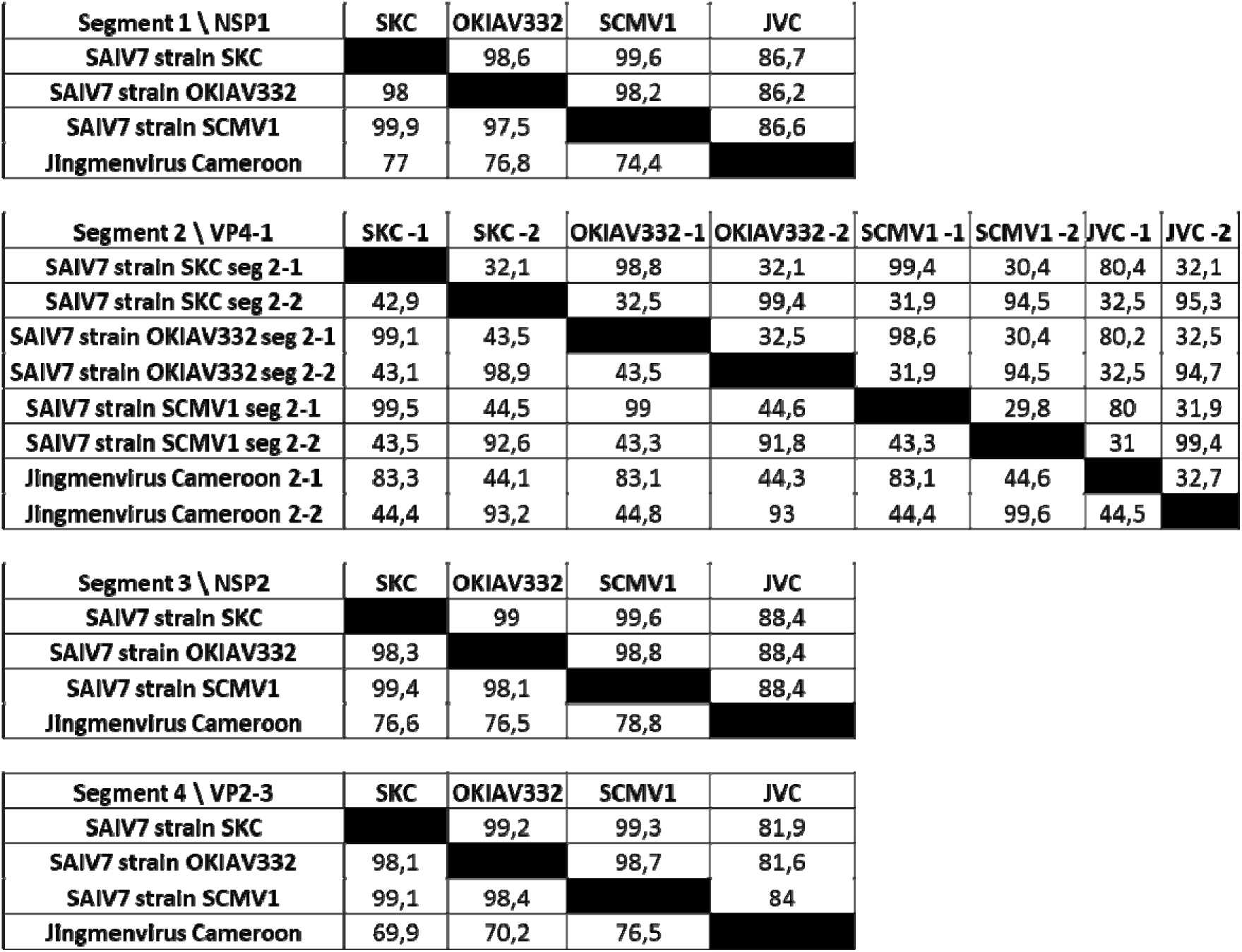
Percentage identity derived from multiple sequence alignment of SAIV7 strains SKC, OKIAV332, SCMV1 and JVC genomic sequences. The bottom left of each table represents nucleotide identity over the whole genomic segment. The top right of each table corresponds to amino acid percentage identity over the corresponding ORF.

These nucleotide sequences (SAIV7 segment 2-2 and JVC segment 2-2) have only one significant match using BLASTn, the published sequence of OKIAV332 seg2-2 (MW314687), but are similar to other jingmenvirus segment 2 sequences using BLASTx (nucleotide query compared to protein databases).

### Genomic fluidity is also found in the tick-associated jingmenvirus clade

Following these discoveries in the insect-associated clade of the jingmenvirus phylogeny, we found a novel jingmenvirus genomic sequence with five segments in dataset SRX3777434 from a single adult female Dysdera bandamae spider body (no legs or pedipalps) from the Canary Islands, Spain in 2015 (Figure 6) (33). This virus was tentatively named Dysdera bandamae jingmenvirus 1 (DBJV1) and its most closely related sequences are the genomic segments of Hainan jingmen-like virus (HJLV), detected in Chinese forest soil metagenome collected in 2018, one of the most divergent sequence in the tick-associated clade of the jingmenvirus phylogeny (Figure 3) (34). DBJV1 genome has two versions of segment 2, one coding for two ORFs (nuORF and VP1; hereafter designated segment 2-1) while the other version does not seem to code for nuORF (hereafter designated segment 2-2). The segment 2-1 nuORF sequence has no match on BLASTx, however we have identified an ORF in the closely related HJLV segment 2 (128–526) that shares 39.6% amino acid identity with DBJV1 nuORF and that was not annotated upon publication.

**Figure 6:**
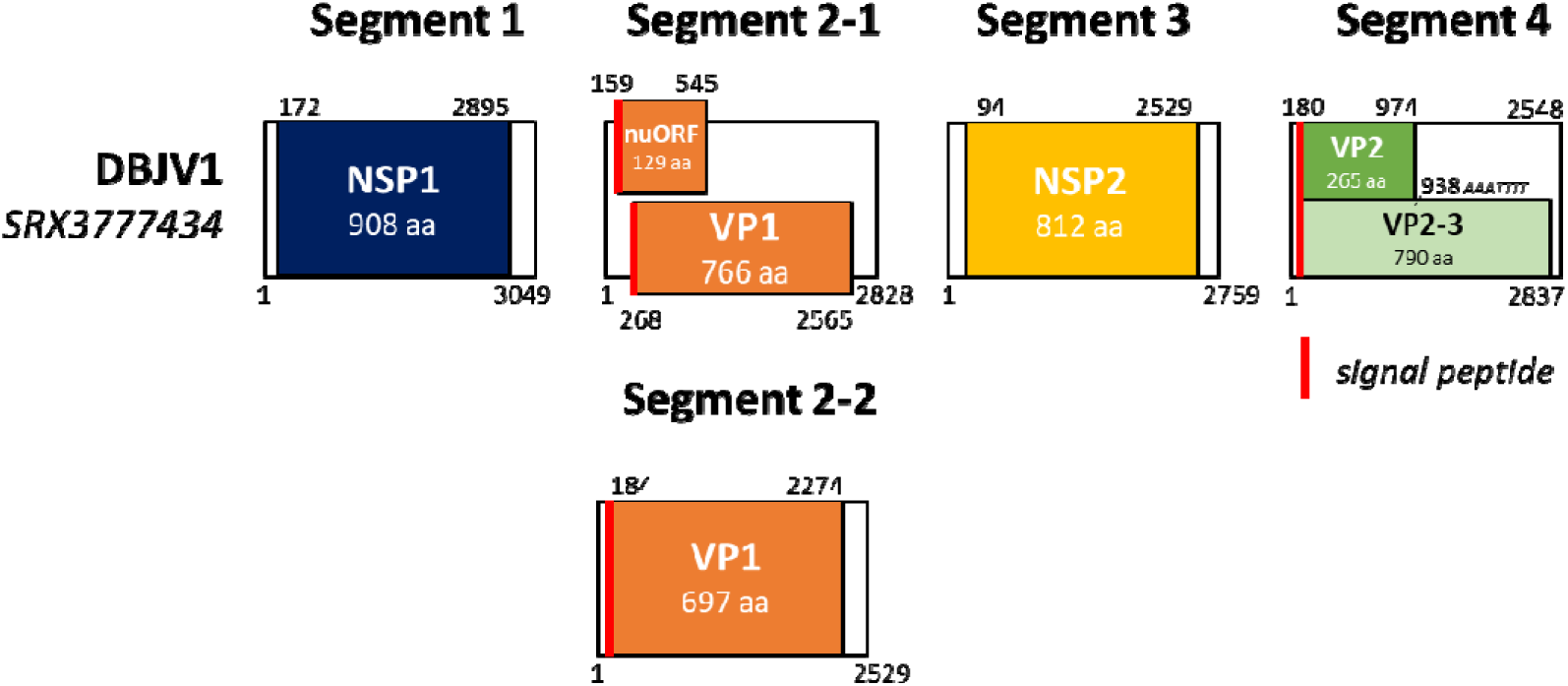
Genomic organisation of Dysdera bandamae jingmenvirus 1 (DBJV1). ORFs are represented by blocks of colour, and translation resulting from a ribosomal frameshift is represented by blocks in pastel colours, with the position of the frameshift and slippery heptanucleotide sequence specified underneath.

Interestingly, the UTRs are more conserved than the coding regions between segment 2-1 (with nuORF and VP1) and segment 2-2 (with VP1 only) as they share 52.3% nucleotide identity (5’ UTR) and 57.1% nucleotide identity (3’ UTR) while the coding regions only share 44.3% nucleotide identity. This was also found for FNQJV1 segment 3-1 (VP1-3 with non-conserved VP3; and VP2) and 3-2 (VP1-3 only) which share 46.3% and 43.4% nucleotide identity over the 5’ and 3’ UTRs respectively while they only share 37.3% nucleotide identity over the coding sequences. This discrepancy was not observed for FNQJV1 and CTJV1 segments 4-1 and 4-2 or for SAIV7 segments 2-1 and 2-2. We were not able to assemble the full UTR sequences of CTJV1 segments 3-1 and 3-2 so the comparison cannot be made for these two segments.

We searched for DBJV1 reads in all other samples of the BioProject containing SRX3777434 and found that none had evidence of DBJV1 sequence, including the datasets containing the metagenome of the legs and pedipalp of the same spider (#2 spider ID CRBA2178). In dataset SRX3777434, the number of reads mapping to the five genomic segment sequences of DBJV1 were as follows: 438 reads for segment 1; 2039 for segment 2-1; 1175 for segment 2-2; 861 for segment 3 and 3440 for segment 4.

### Identification of translation initiation mechanisms

Slippery heptanucleotides have previously been proposed as a mechanism for coding several proteins in a single segment for GCXV, MoCV, JMTV and Alongshan virus. As we analysed the sequences and genomic organisation of the newly discovered viruses, we noticed that most jingmenvirus segments with multiple ORFs contained slippery heptanucleotide motifs that could facilitate a ribosomal frameshift. We therefore systematically analysed genomic sequences of 25 jingmenvirus species for the presence of such motifs.

In the genomes of all mosquito-derived jingmenviruses (GCXV, MoCV, FNQJV1, CTJV1), we identified strictly conserved heptanucleotide motifs in segments 3 (VP1-3 GGAUUUU) and 4 (VP5-6 AAAAAAC) (Table S1). We also identified motifs in all sequences of the more distantly related insect-derived jingmenvirus we analysed (no strict consensus, segment 2 VP4-1; GGUUUUU in most cases, segment 4 VP2-3) and in tick-associated jingmenvirus sequences (segment 2 VP1a-b AAAAAAC; segment 4 VP2-3 GGUUUUU or AAAUUUU) (Table S1).Therefore, ribosomal frameshift is the putative mechanism which initiates translation of multiple ORFs on a single segment and is conserved among all jingmenviruses: tick-associated and insect-associated (including mosquito-derived), and in all versions of homologous segments.

Some remaining ORFs that require alternative translation initiation have no evidence of ribosomal frameshift: VP4/VP5 and VP1/VP2 in mosquito-derived jingmenviruses and nuORF/VP1 in tick-associated jingmenviruses (Figures 1 and 6 for reference). For these ORFs, the mechanism allowing translation initiation also seems conserved and is most likely to be leaky scanning. Indeed, in almost all instances, the first AUG in the segment corresponds to the methionine START of the first ORF to be translated (VP4 or VP1 for mosquito-derived jingmenviruses; nuORF for tick-associated jingmenviruses), while the second AUG corresponds to the methionine START of the second ORF to be translated (VP5 or VP1 for mosquito-derived jingmenviruses; VP1 or VP1a for tick-associated jingmenviruses) (Table S2). For some viruses the methionine START of the second ORF is the second or third instance of AUG after the initial one (Table S2).

Finally, we used the tool DeepIRES to screen for the presence of IRES in the jingmenvirus genomic segments with multiple ORFs and found no instances where the predictions from DeepIRES matched a predicted ORF, suggesting this mechanism for alternative translation initiation might not be relevant to jingmenvirus biology.

The predicted mechanisms we propose here (ribosomal frameshift and leaky scanning) are conserved for all jingmenvirus sequences analysed but need to be confirmed with biological samples, for example using mass spectrometry as it was done to confirm ribosomal frameshifting in GCXV segment 3 (3).

### Structural analysis

We chose the designations segment x-1 and x-2 based on the genomic organisation or on nucleotide percentage identity for sequences when homology to known jingmenvirus sequences (in this case, to GCXV) could be identified. However, for jingmenviruses more distant than the mosquito-derived jingmenviruses are to each other, nucleotide or amino acid sequences comparisons yield low to no homology. This observation applies for the copies of segments from a same virus (Figure 2 and Figure 5). In order to determine whether the segments we identified code for homologous proteins, we generated structural models for the putative structural proteins in the FNQJV1, CTJV1, SAIV7 and GCXV sequences using AlphaFold and performed structural alignments with FATCAT and Chimera X. Using this method, we also attempted to confirm which segments encode similar proteins between the mosquito-derived sequences (GCXV, CTJV1, FNQJV1) and more distantly related SAIV7, as its genomic organization follows a different nomenclature compared to mosquito-derived jingmenviruses, due to low sequence identity in the segments that are not coding for non-structural proteins.

We first generated the structural models of VP1 the only protein in common for GCXV segment 3, FNQJV1 segments 3-1 and 3-2, CTJV1 segments 3-1 and 3-2. The models were superimposed in order to generate a structural alignment. The structures showed statistically significant similarity (Table S3) and conserved motifs were identified (Figure 7). VP1 is organised in three domains, a N-terminal domain composed of three α-helices, a central domain containing three antiparallel β-strands, and a C-terminal domain organised in six β-strands surrounded by five α-helices (Figure 7). We then performed pairwise comparisons between GCXV VP1 and SAIV7 putative structural proteins and found that GCXV VP1 shares statistically significant structure similarities with SAIV7 VP2 (*p* = 1.38 x 10^-4^, Table S3) but is not statistically significantly similar to SAIV7 VP4 (*p* = 0.459 and 0.271 for segment 2-1 and 2-2 respectively). We were therefore able to include SAIV7 VP2 in the multiple structure alignment presented here (Figure 7, Table S3) and identify the three domain-organization, and conserved residues and motifs shared between SAIV7 VP2 and all mosquito-derived jingmenvirus VP1 sequences, including four cysteines involved in two putative disulphide bonds contributing to the structural organization of the N- and C-terminal domains.

**Figure 7:**
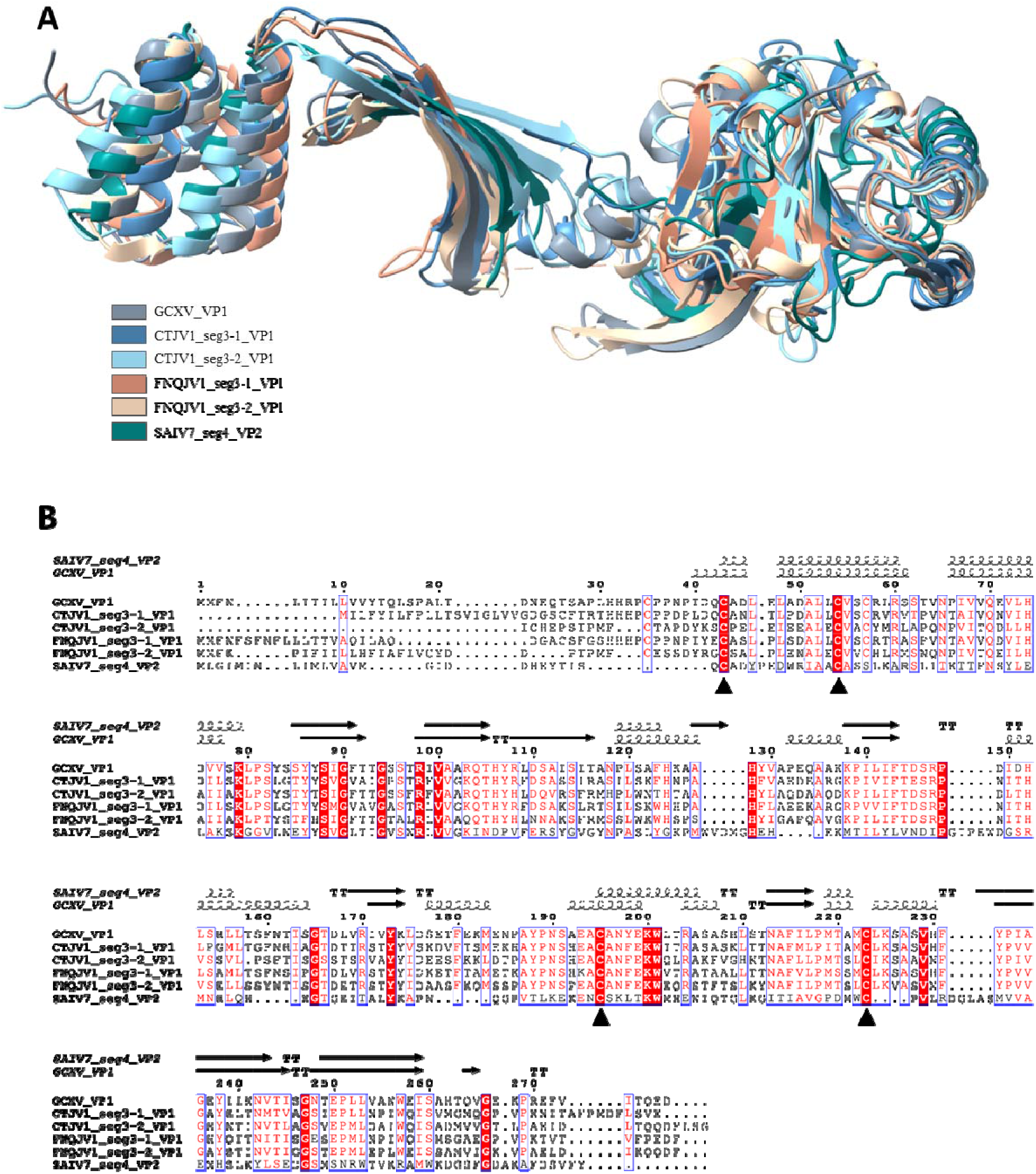
Structural comparison of representative structures of VP1 and bioinformatic analysis. **(A)** GCXV_VP1 (grey), CTJV1_seg3-1_VP1 (beige), CTJV1_seg3-2_VP1 (blue), FNQJV1_seg3-1_VP1 (green), FNQJV1_seg3-2_VP1 (orange) and SAIV7_VP2 (pink) structures were generated using AlphaFold (35). Structures were superimposed using flexible FATCAT (23) and visualized in Chimera X (24). **(B)** A structural alignment was generated by Chimera X and analysed with ESPript. The numbers on top of the alignment indicate the amino acid positions in the GCXV_VP1 sequence. Spirals and arrows above the alignment indicate the position of α-helices and β-strands, based on the AlphaFold models. Cysteines that may be involved in disulfide bridges are indicated by ▴.

Finally, we aligned the structures of VP4 sequences from GCXV segment 4, FNQJV1 segment 4-1 and 4-2, CTJV1 segment 4-1 and 4-2 and found significant similarity for all (Table S3, Figure 8). We also included both versions of VP4 from SAIV7 in the multiple structure alignment and found significant similarity between the seven VP4 structures (Figure 8), all organised in five α-helices.

**Figure 8:**
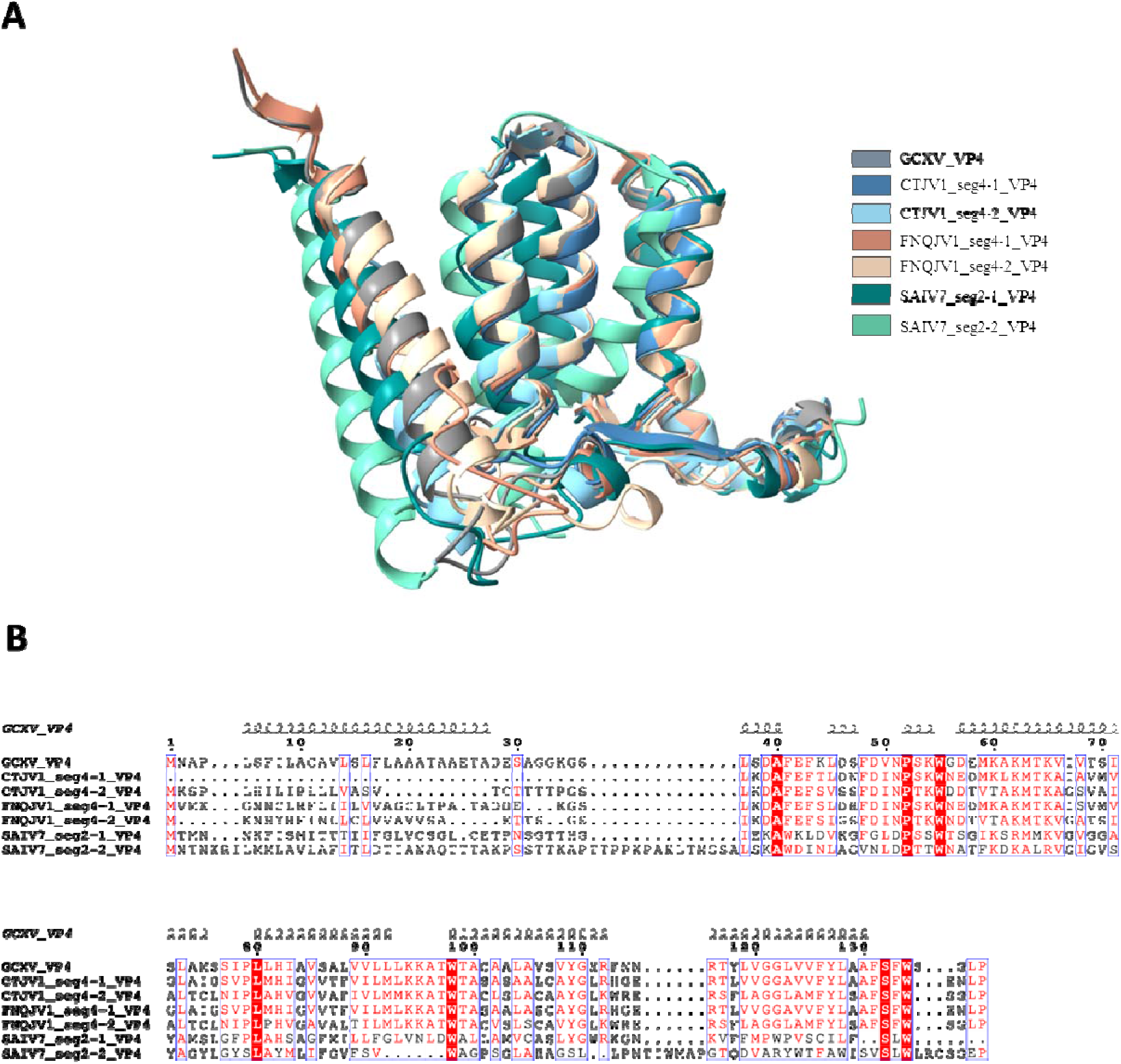
Structural comparison of representative structures of VP4 and bioinformatic analysis. . **(A)** GCXV_VP4 (grey), CTJV1_seg4-1_VP4 (dark blue), CTJV1_seg4-2_VP4 (light blue), FNQJV1_seg4-1_VP4 (dark orange), FNQJV1_seg4-2_VP4 (light orange), SAIV7_seg2-1_VP4 (dark green) and SAIV7_seg2-2_VP4 (light green) structures were generated using AlphaFold (35). Structures were superimposed using flexible FATCAT (23) and visualized in Chimera X (24). **(B)** A structural alignment was generated by Chimera X and analysed with ESPript. The numbers on top of the alignment indicate the amino acid positions in the GCXV_VP4 sequence. Spirals above the alignment indicate the position of α-helices, based on the AlphaFold models.

The structural models of mosquito-derived VP2, VP5-6 and VP7 (GCXV only) generated with AlphaFold were not obtained with sufficient confidence to be included in this study (Table S3, Figure S1).

## Discussion

In this study, we report the discovery of three novel jingmenvirus genomes, which have prompted us to revisit our knowledge of the genomic organisation of jingmenviruses. Until now, jingmenviruses were described as multisegmented viruses with positive single-stranded RNA genome containing 4 segments, with the exception of GCXV, with a genome containing 5 segments. Using a screening approach based on similarity to known jingmenvirus RdRp sequences in public sequencing raw data, we identified that more than four jingmenvirus genomic segments co-occur in different datasets, and that this observation was valid for both the tick-associated and insect-associated clades of the jingmenvirus phylogeny. In all instances, all versions of each segment were always found together in sequencing datasets, including in a dataset derived from a single spider (SRX3777434; DBJV1). In addition to this, we found a single version of the segments coding for the two non-structural proteins NSP1 and NSP2 in each dataset, and homologous versions of other segments. Taken together, these data suggest that the jingmenvirus sequences we found in one dataset are part of a single virus species, and that jingmenviruses could have up to 6 segments.

In addition to discovering new jingmenvirus genomic sequences with more than 4 segments, we uncovered the existence of additional segments in two published jingmenvirus species (SAIV7 and JVC). These newly uncovered segments had remained unidentified in the studies describing these virus species most likely because only four segments were expected to be found in jingmenvirus genomes. It would seem that this is not an isolated occurrence: while this manuscript was in preparation, Tang *et al.* published a new tool to reconstruct segmented virus genomes from metatranscriptomics data: SegVir, and Liu *et al.* published a pre-print article on a tool to identify RNA virus genome segments: SegFinder (36,37). Liu *et al.* conclude that they have identified multiple occurrences of additional genomic segments in virus species with multisegmented and multipartite genomes, compared to the number of segments classically described. This observation is reinforced by the recent identification of jingmen-related sequences in plant-derived samples using the conserved untranslated regions as markers (38). Indeed, Ailanthus jingmen-related virus 1 was identified in the metagenome of the *Ailanthus altissima* tree and was found to have six genomic segments, two related to the jingmenvirus NSP1 and NSP2 respectively, and four sequences with low identity to jingmenvirus sequences, including two segments with ORFs similar to each other (38). These data closely match our findings, and are in agreement with the existence of an evolutionary continuum shared by insect and plant viruses. These publications, taken together with our study, highlight that with the existing knowledge on multipartite viruses, assigning their genomic segments is not straightforward especially when using metagenomics data rather than sequencing an isolate. While we cannot prove without a doubt with the data currently available that the duplicated segments identified here complete the genome of a single virus species, the arguments developed above are in favour of this hypothesis. In order to confirm the number of genomic segments within a single viral genome, and identify the role of each in viral replication, it would be valuable to obtain an isolate of one of these viruses, other than GCXV which does not seem to contain any multi-copy segment (raw sequencing data were unavailable publicly, but the virus isolate reported have 4 or 5 segments), and to develop associated molecular and infections tools.

Indeed, it remains to be seen whether the homologous versions of the segments are necessary for infection and/or replication or if they confer some sort of advantage for replication fitness. First, it is notable that GCXV VP7 homologs were not found, in any data set. The function of this protein/segment and the reason for its presence or absence in some isolates remains a mystery, especially considering that no fitness advantage to the presence of this segment was identified *in vitro* or *in vivo* to date (3,6). Then, in the sequences identified, some homologous segments had ORFs that were absent (nuORF in DBJV1 segment 2-2; VP2 in FNQJV1 and CTJV1 segment 3-2), truncated or poorly conserved protein sequences (VP3 in FNQJV1 and CTJV1 segment 3-1). In other instances, we did not identify organisational differences between the ORFs of the two homologous versions of segment (SAIV7 segments 2-1 and 2-2; FNQJV1 and CTJV1 segments 4-1 and 4-2). This discrepancy could suggest different evolutive processes that drove jingmenvirus evolution (Figure 8). It should be noted that for the ORFs that were conserved in all versions, we were able to find statistically significant structural homology between the proteins they encoded (mosquito-derived VP1 and VP4 for example), despite the sequence divergence observed. The biological relevance of this structural redundancy is not clear at this stage, as structural conservation is not necessary for functional conservation as exemplified by the Rossmann fold, a prevalent structural motif that can accommodate a wide range of functions (39).

Given the structural and organisational (conservation of slippery sequences, order and number of ORFs) conservation of the segments and related viral proteins, it would be tempting to classify these homologous segments as paralogs, resulting from a duplication of the parental segment, followed by a drift of their sequence. However, multipartite viruses seem to be more tolerant than their monopartite counterparts to fluidity in their genomic organisation and to evolutive processes that facilitate functional gain by different mechanisms, including duplication recombination, reassortment or gene acquisition (5). As mentioned in the introduction, multipartite virus particles encapsidate less fragments than the total number of genomic segments, multiple particles are required to initiate the replication cycle and they are extremely rare in animals (5). The genome plasticity observed here for jingmenviruses would be in line with a multipartite organisation. Indeed, the fluid genomic organisation could be explained by packaging of different combinations of segments leading to a reshuffling of genomic diversity and organisation, with gain or loss of genomic material and gain, loss or overlap of function (Figure 8). The segment duplication observed here could result from different mechanisms, for example a virus with an initial infectious unit of 4 segments acquiring an extra version of a gene from another virus species during co-infection, therefore becoming a virus with an infectious unit of 5 segments - as opposed to segments reassortment, which would not result in a change in the number of segments in the infectious unit. Another possible mechanism for gene duplication in multipartite jingmenviruses is dynamic evolution by several rounds of adaptive pressure selection on multiple quasi-species at once in one virus species, once again increasing the number of segments in the infectious unit by one, by adding a homolog of an existing gene to the infectious unit. These mechanisms would be facilitated by the putative multipartite organisation, with packaging of different combinations of segments in different virus particles as mentioned above, and therefore requiring multiple particles to obtain an infectious unit.

**Figure 9:**
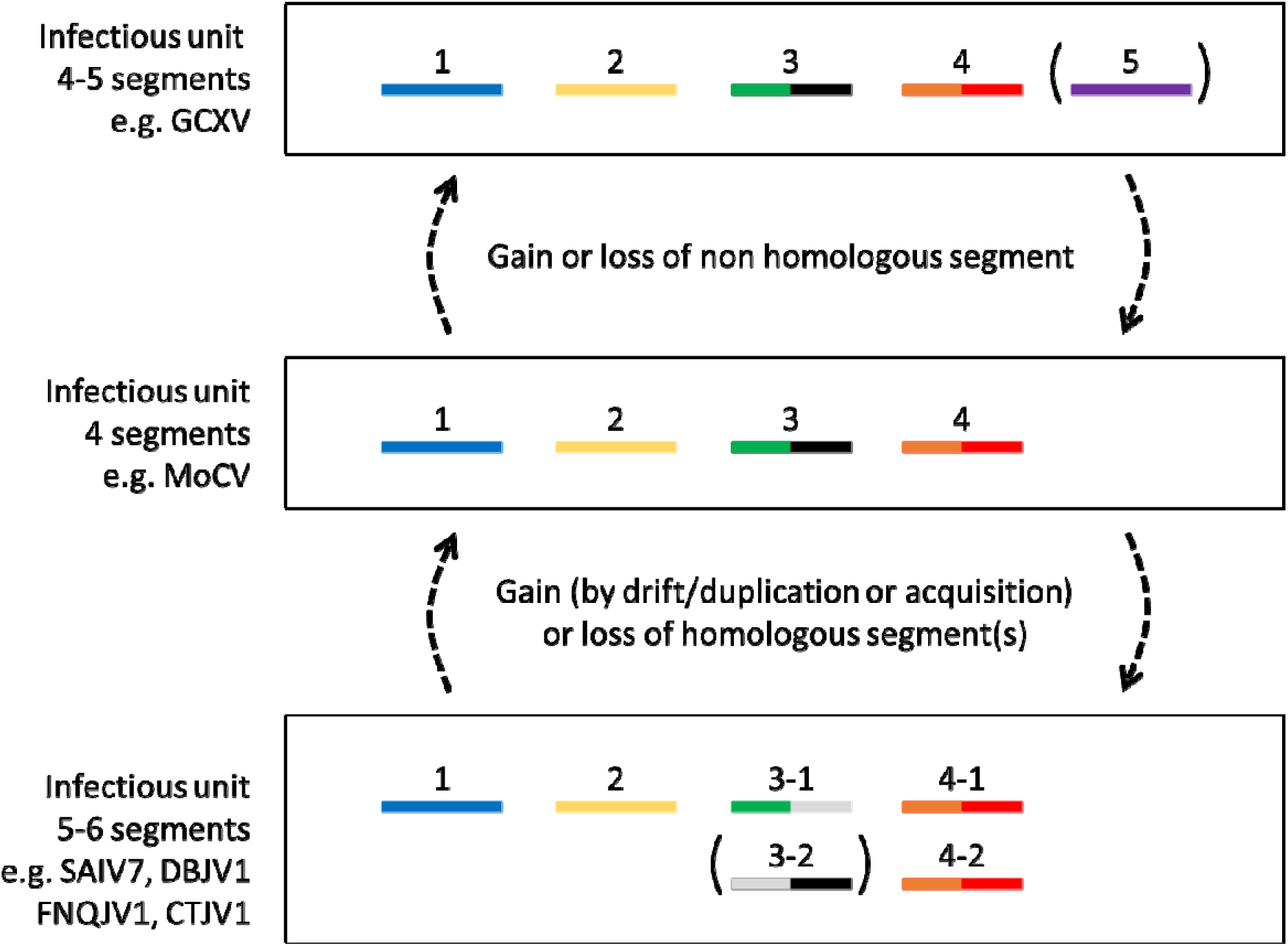
Fluidity of the jingmenvirus genomic organisation. Jingmenviruses contain 4 to 6 homologous or non-homologous genomic segments per infectious unit. These homologous versions may share the same organisation (ORFs, size) or may have complementary putative functions with ORFs lacking in one version (3-1 vs 3-2) but not the other (4-1 vs 4-2), for example. Different ORFs are represented in different colours, with grey representing missing or non-conserved ORFs.

Moreover, for all sequences identified here, we observed variations in the sequencing depth between segments and between datasets, which suggests a variation in the quantity of RNA of each segment in the different biological samples. This could be linked to the notion of genomic formula for multipartite viruses, as it has been observed that different segments of the multipartite genome might vary in orders of magnitude with relative frequencies that seem to be host-specific (40). With these observations and the conclusions from Ladner et al. on GCXV following a virus dilution and low multiplicity of infection experiment (3) raises the hypothesis that the presence of homologous segments could provide a selective advantage by increasing the host spectrum of jingmenviruses.

This study has highlighted the need for a simpler nomenclature for jingmenvirus genomic organisation. The discrepancies between segment numbers and ORF designations have and will lead to confusion when comparing tick-associated (for example JMTV, DBJV1), insect-derived (for example SAIV7) and mosquito-derived (GCXV, MoCV, FNQJV1, CTJV1) jingmenvirus genomes. Here we confirm a structural homology between proteins from mosquito-derived jingmenvirus genomes and insect-derived jingmenvirus genomes which have a different nomenclature: VP1 from mosquito-derived jingmenvirus segment 3 and VP2 from insect-derived jingmenvirus segment 4; and VP4 from mosquito-derived jingmenvirus segment 4 and VP4 from insect-derived jingmenvirus segment 2. The structural comparisons yielded results with a much higher confidence than those obtained by sequence comparisons, as the percentage identities obtained by multiple sequence alignment were similar to those that can be obtained when comparing two unrelated sequences (≤ 30%). Continuing to confirm homology between ORFs in different clades of the jingmenvirus phylogeny would help simplify the nomenclature.

Considering the fluid nature of the jingmenvirus genome highlighted in this study, and the high sequence divergence observed between sequences in different phylogenetic clades, the simplest nomenclature would be to have segment 1 carrying the RdRp and methyltransferase functional domains; segment 2 carrying the serine protease and helicase domains; segment 3 carrying the putative envelope; segments 4,5,6… for additional segments with unknown or untested functions, ranked according to size, as has historically been the case; a numbered suffix for any obvious homolog (3-1, 3-2 for example). Search of distant homology by protein structures alignments in addition to nucleotide or amino acid sequences alignment to attribute the segment number would contribute to rationalizing the nomenclature. ORFs with ribosomal frameshifts could be named ORF1a and ORF1ab for the version coding for the shorter or longer protein respectively, coding for VP1 and VP1-3, rather than being named VP1 and VP3 (for example). Once the function of these ORFs encoding putative structural proteins is confirmed, the product of these ORFs could be renamed with that function (envelope for example) rather than “VP” followed by a number, that might differ between jingmenviruses from different clades. Implementing these changes could result in challenging circumstances if already published records were not corrected.

In addition to these proposed modifications in nomenclature, special care should be taken when annotating the ORFs in a new jingmenvirus genome. Indeed, based on sequence analyses, we propose different mechanisms for the alternative translation initiation in jingmenvirus segments with multiple ORFs: ribosomal frameshifts and leaky scanning, depending on the segment and ORF (13). In particular, the ribosomal frameshifts lead to updated ORF annotations for a number of segments in a number of jingmenvirus species (for example VP2-3 rather than VP2 and a separate VP3). Again, biological verification would be needed to confirm the relevance of these proposed molecular mechanisms, for example the same way that GCXV ribosomal frameshifts were confirmed: with peptides dependent on the frameshift detected in virus samples by mass spectrometry (3).

Overall, particular care should be taken when publishing and depositing new jingmenvirus sequences, for all the reasons discussed above: considering the presence of homologous segments when metagenomic data are analysed will help to find “hidden” segments and ORFs on segments with multiple ORFs should be able to be translated following a molecular mechanism to be biologically relevant.

## Conclusion

With the discovery of novel jingmenvirus-related sequences, we have been able to put forward new evidence of the multipartite nature of jingmenviruses, and of the evolutionary role this organisation may play. With these new considerations, a harmonized nomenclature should be put in place whenever possible, aiming at clarifying, correcting and simplifying the formal genomic organisation of jingmenviruses.

## Acknowledgements

The authors would like to thank Jean-Marc Haselhoff, Morgan Seston and Aurélien Destruel for helpful technical help; as well as Dr Michael Berg and Dr Gregory Orf for uploading the raw sequencing data from (32) to NCBI’s SRA database when we requested it.

## Data availability

All newly described sequences were deposited in Genbank under accession numbers XXXXXXX-XXXXXXXX (submitted).

Funding

This work was supported by the European Virus Archive GLOBAL (EVA-GLOBAL) project that has received funding from the European Union’s Horizon 2020 Research and Innovation Program under grant agreement no. 871029.

## References

1. Colmant AMG, Charrel RN, Coutard B. Jingmenviruses: Ubiquitous, understudied, segmented flavi-like viruses. Front Microbiol. 2022;13:997058.

2. Qin XC, Shi M, Tian JH, Lin XD, Gao DY, He JR, et al. A tick-borne segmented RNA virus contains genome segments derived from unsegmented viral ancestors. Proc Natl Acad Sci U S A. 2014 May 6;111(18):6744–9.

3. Ladner JT, Wiley MR, Beitzel B, Auguste AJ, Dupuis AP, Lindquist ME, et al. A Multicomponent Animal Virus Isolated from Mosquitoes. Cell Host Microbe. 2016 Sep 14;20(3):357–67.

4. Pauvolid-Corrêa A, Solberg O, Couto-Lima D, Nogueira RM, Langevin S, Komar N. Novel Viruses Isolated from Mosquitoes in Pantanal, Brazil. Genome Announc. 2016 Nov 3;4(6):e01195–16.

5. Michalakis Y, Blanc S. The Curious Strategy of Multipartite Viruses. Annu Rev Virol. 2020 Sep 29;7(1):203–18.

6. Chen RY, Zhao T, Guo JJ, Zhu F, Zhang NN, Li XF, et al. The infection kinetics and transmission potential of two Guaico Culex viruses in Culex quinquefasciatus mosquitoes. Virol Sin. 2024 Mar 8;S1995–820X(24)00028-2.

7. Wang X, Jing X, Shi J, Liu Q, Shen S, Cheung PPH, et al. A jingmenvirus RNA-dependent RNA polymerase structurally resembles the flavivirus counterpart but with different features at the initiation phase. Nucleic Acids Res. 2024 Jan 31;gkae042.

8. Chen H, Lin S, Yang F, Chen Z, Guo L, Yang J, et al. Structural and functional basis of low-affinity SAM/SAH-binding in the conserved MTase of the multi-segmented Alongshan virus distantly related to canonical unsegmented flaviviruses. PLoS Pathog. 2023 Oct;19(10):e1011694.

9. Mifsud JCO, Lytras S, Oliver MR, Toon K, Costa VA, Holmes EC, et al. Mapping glycoprotein structure reveals Flaviviridae evolutionary history. Nature. 2024 Sep;633(8030):695–703.

10. Gao X, Zhu K, Wojdyla JA, Chen P, Qin B, Li Z, et al. Crystal structure of the NS3-like helicase from Alongshan virus. IUCrJ. 2020 May 1;7(Pt 3):375–82.

11. Zhang XY, Shu T, Wang X, Xu J, Qiu Y, Zhou X. Guaico Culex virus NSP2 has RNA helicase and chaperoning activities. J Gen Virol. 2021 Apr;102(4).

12. Amoa-Bosompem M, Kobayashi D, Murota K, Faizah AN, Itokawa K, Fujita R, et al. Entomological Assessment of the Status and Risk of Mosquito-borne Arboviral Transmission in Ghana. Viruses. 2020 Jan 27;12(2):E147.

13. Sorokin II, Vassilenko KS, Terenin IM, Kalinina NO, Agol VI, Dmitriev SE. Non-Canonical Translation Initiation Mechanisms Employed by Eukaryotic Viral mRNAs. Biochemistry (Mosc). 2021;86(9):1060–94.

14. Shi M, Lin XD, Vasilakis N, Tian JH, Li CX, Chen LJ, et al. Divergent Viruses Discovered in Arthropods and Vertebrates Revise the Evolutionary History of the Flaviviridae and Related Viruses. J Virol. 2016 Jan 15;90(2):659–69.

15. Kholodilov IS, Litov AG, Klimentov AS, Belova OA, Polienko AE, Nikitin NA, et al. Isolation and Characterisation of Alongshan Virus in Russia. Viruses. 2020 Mar 26;12(4):E362.

16. Edgar RC, Taylor J, Lin V, Altman T, Barbera P, Meleshko D, et al. Petabase-scale sequence alignment catalyses viral discovery. Nature. 2022 Feb;602(7895):142–7.

17. Vandegrift KJ, Kumar A, Sharma H, Murthy S, Kramer LD, Ostfeld R, et al. Presence of Segmented Flavivirus Infections in North America. Emerg Infect Dis. 2020 Aug;26(8):1810–7.

18. Lefort V, Longueville JE, Gascuel O. SMS: Smart Model Selection in PhyML. Mol Biol Evol. 2017 Sep 1;34(9):2422–4.

19. McNair K, Salamon P, Edwards RA, Segall AM. PRFect: a tool to predict programmed ribosomal frameshifts in prokaryotic and viral genomes. BMC Bioinformatics. 2024 Feb 22;25(1):82.

20. Andronescu M, Aguirre-Hernández R, Condon A, Hoos HH. RNAsoft: a suite of RNA secondary structure prediction and design software tools. Nucleic Acids Res. 2003 Jul 1;31(13):3416–22.

21. Zhao J, Chen Z, Zhang M, Zou L, He S, Liu J, et al. DeepIRES: a hybrid deep learning model for accurate identification of internal ribosome entry sites in cellular and viral mRNAs. Brief Bioinform. 2024 Jul 25;25(5):bbae439.

22. Mirdita M, Schütze K, Moriwaki Y, Heo L, Ovchinnikov S, Steinegger M. ColabFold: making protein folding accessible to all. Nat Methods. 2022 Jun;19(6):679–82.

23. Li Z, Jaroszewski L, Iyer M, Sedova M, Godzik A. FATCAT 2.0: towards a better understanding of the structural diversity of proteins. Nucleic Acids Res. 2020 Jul 2;48(W1):W60–4.

24. Pettersen EF, Goddard TD, Huang CC, Couch GS, Greenblatt DM, Meng EC, et al. UCSF Chimera--a visualization system for exploratory research and analysis. J Comput Chem. 2004 Oct;25(13):1605–12.

25. Robert X, Gouet P. Deciphering key features in protein structures with the new ENDscript server. Nucleic Acids Res. 2014 Jul;42(Web Server issue):W320–324.

26. Ramírez AL, Colmant AMG, Warrilow D, Huang B, Pyke AT, McMahon JL, et al. Metagenomic Analysis of the Virome of Mosquito Excreta. mSphere. 2020 Sep 9;5(5):e00587–20.

27. Xiao P, Han J, Zhang Y, Li C, Guo X, Wen S, et al. Metagenomic Analysis of Flaviviridae in Mosquito Viromes Isolated From Yunnan Province in China Reveals Genes From Dengue and Zika Viruses. Front Cell Infect Microbiol. 2018;8:359.

28. Li CX, Shi M, Tian JH, Lin XD, Kang YJ, Chen LJ, et al. Unprecedented genomic diversity of RNA viruses in arthropods reveals the ancestry of negative-sense RNA viruses. Elife. 2015 Jan 29;4:e05378.

29. Paraskevopoulou S, Käfer S, Zirkel F, Donath A, Petersen M, Liu S, et al. Viromics of extant insect orders unveil the evolution of the flavi-like superfamily. Virus Evol. 2021 Jan;7(1):veab030.

30. Zhao M, Yue C, Yang Z, Li Y, Zhang D, Zhang J, et al. Viral metagenomics unveiled extensive communications of viruses within giant pandas and their associated organisms in the same ecosystem. Sci Total Environ. 2022 May 10;820:153317.

31. He WT, Hou X, Zhao J, Sun J, He H, Si W, et al. Virome characterization of game animals in China reveals a spectrum of emerging pathogens. Cell. 2022 Mar 31;185(7):1117–1129.e8.

32. Orf GS, Olivo A, Harris B, Weiss SL, Achari A, Yu G, et al. Metagenomic Detection of Divergent Insect- and Bat-Associated Viruses in Plasma from Two African Individuals Enrolled in Blood-Borne Surveillance. Viruses. 2023 Apr 21;15(4):1022.

33. Vizueta J, Macías-Hernández N, Arnedo MA, Rozas J, Sánchez-Gracia A. Chance and predictability in evolution: The genomic basis of convergent dietary specializations in an adaptive radiation. Mol Ecol. 2019 Sep;28(17):4028–45.

34. Chen YM, Sadiq S, Tian JH, Chen X, Lin XD, Shen JJ, et al. RNA viromes from terrestrial sites across China expand environmental viral diversity. Nat Microbiol. 2022 Aug;7(8):1312–23.

35. Jumper J, Evans R, Pritzel A, Green T, Figurnov M, Ronneberger O, et al. Highly accurate protein structure prediction with AlphaFold. Nature. 2021 Aug;596(7873):583–9.

36. Tang X, Shang J, Chen G, Chan KHK, Shi M, Sun Y. SegVir: Reconstruction of Complete Segmented RNA Viral Genomes from Metatranscriptomes. Mol Biol Evol. 2024 Aug 2;41(8):msae171.

37. Liu X, Kong J, Shan Y, Yang Z, Miao J, Pan Y, et al. SegFinder: an automated tool for identifying RNA virus genome segments through co-occurrence in multiple sequenced samples. bioRxiv; 2024. p. 2024.08.19.608591

38. Zhang S, Yang C, Qiu Y, Liao R, Xuan Z, Ren F, et al. Conserved untranslated regions of multipartite viruses: Natural markers of novel viral genomic components and tags of viral evolution. Virus Evol. 2024;10(1):veae004.

39. Medvedev KE, Kinch LN, Dustin Schaeffer R, Pei J, Grishin NV. A Fifth of the Protein World: Rossmann-like Proteins as an Evolutionarily Successful Structural unit. J Mol Biol. 2021 Feb 19;433(4):166788.

40. Lucía-Sanz A, Manrubia S. Multipartite viruses: adaptive trick or evolutionary treat? npj Syst Biol Appl. 2017 Nov 9;3(1):1–11.

