## Supplementary figures for "The fluid genomic organisation of jingmenviruses"

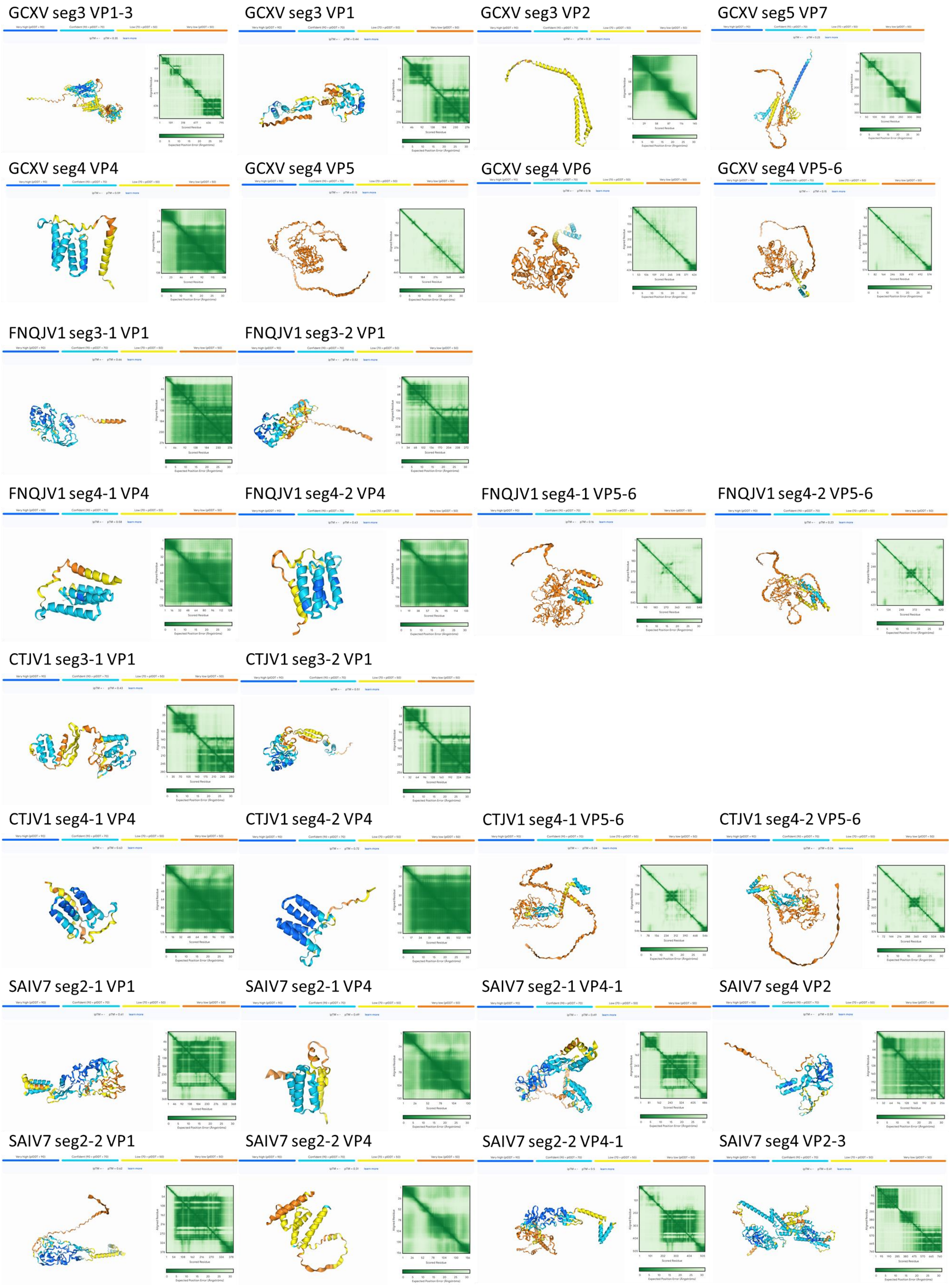

Figure S1 Putative structural protein structures, generated with AlphaFold.

**Table S1: Heptanucleotide slippery motifs identified in jingmenvirus sequences**

| Virus (Accession) | Segment # (ORFs) | Position | Heptanucleotide identified |
|---|---|---|---|
| TAKV (LC628181) | 2 (VP1a-b) | 1399-1405 | AAAAAAC |
| ALSV (MH158416) | 2 (VP1a-b) | 1431-1437 | AAAAAAC* (1) |
| YGTV (MH688530) | 2 (VP1a-b) | 1417-1423 | AAAAAAC |
| XTJV1 (MZ244285) | 2 (VP1a-b) | 1360-1366 | AAAAAAC |
| GCXV (KM461669) | 4 (VP5-6) | 739-745 | AAAAAAC* (2) |
| MoCV (LC505055) | 4 (VP5-6) | 721-727 | AAAAAAC* (3) |
| CTJV1 (accession) | 4-1 (VP5-6) | 644-650 | AAAAAAC |
| CTJV1 (accession) | 4-2(VP5-6) | 532-538 | AAAAAAC |
| FNQJV1 (accession) | 4-1 (VP5-6) | 846-852 | AAAAAAC |
| FNQJV1 (accession) | 4-2 (VP5-6) | 595-601 | AAAAAAC |
| IFJV1 (OM869460) | 2 (VP4-1) | 290-296 | TTTGGGC |
| WHCV (KR902710) | 2 (VP4-1) | 378-385 | UAAUUUU |
| WHFV (KR902714) | 2 (VP4-1) | 437-443 | UUUUUUA |
| SAIV7 (KR902718) | 2-1 (VP4-1) | 442-448 | GGUUUUU |
| WHAV1 (KR902722) | 2 (VP4-1) | 397-403 | AAAAACC |
| WHAV2 (KR902726) | 2 (VP4-1) | 311-317 | GGCUUUU |
| SBTV2 (MW023855) | 2 (VP4-1) | 445-451 | GGUUUUU |
| SBTV3 (MW033629) | 2 (VP4-1) | 408-414 | AAAAAAG |
| OKIAV339 (MW208801) | 2 (VP4-1) | 415-421 | AAUUUUU |
| OKIAV337 (MW314691) | 2 (VP4-1) | 444-450 | UUUUUUA |
| JMTV (KJ001582) | 4 (VP2-3) | 898-904 | GGUUUUU* (4) |
| TAKV (LC628183) | 4 (VP2-3) | 850-856 | GGUUUUU |
| ALSV (MH158418) | 4 (VP2-3) | 849-855 | GGUUUUU* (1) |
| YGTV (MH688532) | 4 (VP2-3) | 872-878 | GGUUUUU |
| GJLV (MW896896) | 4 (VP2-3) | 920-926 | GGUUUUU |
| XTJV1 (MZ244284) | 4 (VP2-3) | 861-867 | GGUUUUU |
| DBJV1 (accession) | 4 (VP2-3) | 938-944 | AAAUUUU |
| HJLV (MW896923) | 4 (VP2-3) | 855-861 | AAAUUUU |
| GCXV (KM461668) | 3 (VP1-3) | 950-957 | GGAUUUU* (2) |
| MoCV (LC505054) | 3 (VP1-3) | 933-939 | GGAUUUU* (3) |
| CTJV1 (accession) | 3-1 (VP1-3) | 839-845 | GGAUUUU |
| CTJV1 (accession) | 3-2 (VP1-3) | 757-763 | GGAUUUU |
| FNQJV1 (accession) | 3-1 (VP1-3) | 1010-1016 | GGAUUUU |
| FNQJV1 (accession) | 3-2 (VP1-3) | 885-891 | GGAUUUU |
| IFJV1 (OM869462) | 4 (VP2-3) | 836-842 | GGAUUUU |
| WHCV (KR902712) | 4 (VP2-3) | 828-834 | GGAUUUU* (4) |
| WHFV (KR902716) | 4 (VP2-3) | 855-861 | UUUUUUA |
| SAIV7 (KR902720) | 4 (VP2-3) | 850-856 | GGUUUUU |
| WHAV1 (KR902724) | 4 (VP2-3) | 906-912 | GGUUUUU |
| WHAV2 (KR902728) | 4 (VP2-3) | 887-893 | GGUUUUU |
| CarV (LC552038) | 4 (VP2-3) | 872-878 | GGUUUUU |
| SBTV1 (MW023853) | 4 (VP2-3) | 847-853 | CCCGGGG |
| SBTV2 (MW023857) | 4 (VP2-3) | 877-883 | GGUUUUU |
| SBTV3 (MW033631) | 4 (VP2-3) | 894-900 | GGUUUUU |
| SBTV4 (MW033627) | 4 (VP2-3) | 883-890 | GGUUUUU |
| SBTV5 (MW033632) | 4 (VP2-3) | 880-886 | GGUUUUU |
| OKIAV339 (MW208806) | 4 (VP2-3) | 928-934 | GGUUUUU |
| OKIAV337 (MW314693) | 4 (VP2-3) | 892-898 | GGUUUUU |
| CCJV (MZ771211) | 4 (VP2-3) | 910-916 | UUCUUUU |

JMTV: Jingmen tick virus; GCXV: Guaico Culex virus; WHCV: Wuhan cricket virus; WHFV: Wuhan flea virus; SAIV7: Shuangao insect virus 7; WHAV: Wuhan aphid virus 1 and 2; MoCV: Mole Culex virus; CarV: Carajing virus; TAKV: Takachi virus; ALSV: Alongshan virus; YGTV: Yanggou tick virus; PLJV: Pteropus lylei jingmenvirus; SBTV: Soybena thrip virus 1, 2, 3, 4 and 5; OKIAV339: Neuropteran jingmen-related virus; OKIAV337: Trichopteran jingmen-related virus; GJLV: Guangdong jingmen-like virus; HJLV: Hainan jingmen-like virus; XTV: Xinjiang tick virus; CCJV: Culicoides circumscriptus jingmenvirus. *Already identified in previously published data.

(1) Kholodilov IS, Litov AG, Klimentov AS, Belova OA, Polienko AE, Nikitin NA, et al. Isolation and Characterisation of Alongshan Virus in Russia. Viruses. 2020 Mar 26;12(4):E362.

(2) Ladner JT, Wiley MR, Beitzel B, Auguste AJ, Dupuis AP, Lindquist ME, et al. A Multicomponent Animal Virus Isolated from Mosquitoes. Cell Host Microbe. 2016 Sep 14;20(3):357–67.

(3) Amoa-Bosompem M, Kobayashi D, Murota K, Faizah AN, Itokawa K, Fujita R, et al. Entomological Assessment of the Status and Risk of Mosquito-borne Arboviral Transmission in Ghana. Viruses. 2020 Jan 27;12(2):E147.

(4) Shi M, Lin XD, Vasilakis N, Tian JH, Li CX, Chen LJ, et al. Divergent Viruses Discovered in Arthropods and Vertebrates Revise the Evolutionary History of the Flaviviridae and Related Viruses. J Virol. 2016 Jan 15;90(2):659–69.

**Table S2: Sequence between the AUG methionine start sequences of the first and second ORFs in polycistronic jingmenvirus genomic segments with no evidence of ribosomal frameshift.**

| Virus (accession) | Segment | Sequence |
|---|---|---|
| JMTV (KJ001580) | 2 (nuORF/VP1) | **AUG**GCAA**AUG** |
| ALSV (MH158415) | 2 (nuORF/VP1a) | **AUG**GCCAGUCAAACAAUUUCAGACACGAUCAACA**AUG** |
| TAKV (LC628181) | 2 (nuORF/VP1a) | **AUG**GCCAACAGAACCAUCUCGGAAUCAAUCAACU**AUG** |
| YGTV (MH688530) | 2 (nuORF/VP1a) | **AUG**GCUGGUACCAUCUCAGACACCAUCAACACUGGUGCCCAGCAAGUCAAGA**AUG** |
| GJLV (MW896894) | 2 (nuORF/VP1) | **AUG**UCCA**AUG** |
| HJLV (MW896921) | 2 (nuORF/VP1) | **AUG**AAACUGUUCAUUCUUGCCCUAAUAGUCGCGCAGGUUGUCUUCGCCACGGCGCAGG<br>UGACUCCCAAACCCACACCAUCCG**AUG** |
| XTJV1 (MZ244285) | 2 (nuORF/VP1a) | **AUG**GCAGGAAAGACCAUCUCGGACACCAUCAACACCGGGGCCCAGCAGGUCAAGACUGCA<br>UUGGACAAGGUGUUGGGUACUAACUCCUUUAACAUUCUUCCUCUUUUCAUCCUUGCUG<br>CAGC*AUG* GUUCUUCACGGGAGACAUCUACCUCAUUGUCUUCAGCGCCAUCGCCGCUGU<br>GGGACCUGCUGCUCGGGCGGAAUCGGCGCUCGUCCCCUUAUUGGUUGCCGUCCUCAUU<br>CGGAGAAACAGUCGCUCGAUUUCAAGAGCGGCUCUCGCGGUCGCAGCGGCUUGCU**AUG** |
| DBJV1 (accession) | 2 (nuORF/VP1) | **AUG**UCUU*AUG* GACACUUG*AUG* GUUGUUUUGCUUUUGAGCAUAGUCAUCCAGGCUGAA<br>GCAACAAACACCACAGACACUUUCACAGAGGCUGUUACAAAGGCCAUUGAAG**AUG** |
| GCXV (KM461668) | 3 (VP1/VP2) | **AUG***AUG* UUUAAUCUCAUCACCAUCUUGCUGGUUGUUUACACCCAGCUAUCCCCUGCCC<br>UGACCGACA**AUG** |
| MoCV (LC505054) | 3 (VP1/VP2) | **AUG***AUG* UUCAAUCUCAUCACCGUCCUGCUGGUUGUUUACACCCAGCUCCCAUCCGCUC<br>UCACCGACA**AUG** |
| FNQJV1 (accession) | 3-1 (VP1/VP2) | **AUG***AUG* UUUAAUUUCUCAUUUAAUUUUCUUCUCUUAACCACCGUUGCACAAAUUUUAG<br>CACAAG**AUG** |
| CTJV1 (accession) | 3-1 (VP1/VP2) | **AUG**AUAUUAUUUUAUAUAUUGUUUCCUCUCUUGACCAGCGUCAUCGGACUAGUUGUUG<br>GCG**AUG** |
| GCXV (KM461669) | 4 (VP4/VP5) | **AUG**AACGCCCCUCUCUCCUUUAUCCUAGCUUGCGCCGUCCUCUCCCUCUUUCUUGCUGC<br>UGCCACCGCCGCGGAAACUGCUGACGAGAGCGCCGGAGGAAAAGGCUCCCUUUCGGACG<br>CCUUCGAGUUUAAACUCGAUUCCUUCGACGUCAACCCAUCAAA**AUG** |
| MoCV (LC505055) | 4 (VP4/VP5) | **AUG**A*AUG* CUCCUGUCUCCCUAAUCCUAGCUUGCACCGUCCUCCUGACAUUUUCUGCAG<br>CAUUUGCCGCGGAAGAGGAGGCGCCAGCUGACUCCAAGGGCUCGCUUAGCGACGCCUUU<br>GAGUUUAAGCUCGACUCCUUCGACGUCAACCCGGCUAA**AUG** |
| FNQJV1 (accession) | 4-1 (VP4/VP5) | **AUG**GUU*AUG* AAAGGAAACAAUUGUCUGCGCUUCCUAAUCAUCCUUGUUGUUGCAGGUU<br>GUUUAACUCCUGCCACUGCUGACGACGAAAAAGGUUCGCUAAAAGACGCUUUCGAAUUU<br>UCCCUUGAUCACUUUGACAUAAAUCCAUCAAA**AUG** |
| FNQJV1 (accession) | 4-2 (VP4/VP5) | **AUG**AAAAACCACUAUCACUUUAUUAACCUCCUCUGUUUGGUCGUCGCUGUAGUUUCAGC<br>UGAGACCACUUCUGGCUCCAUCAAGGACGCUUUUGAAUUCAGCAUUGGUUCCUUCGACA<br>UCAAUCCCACGAA**AUG** |
| CTJV1 (accession) | 4-1 (VP4/VP5) | (partial) |
| CTJV1 (accession) | 4-2 (VP4/VP5) | **AUG**AAAUCCCCUCUGCACAUUCUAAUCCCCUUGCUCUUGGUCGCAAGCGUGACUUGUAC<br>CACCACCACUCCUGGUUCGCUUAAAGACGCCUUUGAAUUUAGUGUCAGCUCCUUUGACA<br>UCAACCCGACAAA**AUG** |

**AUG**: start of first ORF
**AUG**: start of second ORF
*AUG* : additional AUG

**Table S3: Statistical relevance of alignments between putative structural proteins of GCXV, FNQJV1, CTJV1 and SAIV7 represented by the p-value obtained with FATCAT. First table: Top right: comparisons between GCXV VP1, FNQJV1 VP1 from segments 3-1 and 3-2, CTJV1 VP1 from segments 3-1 and 3-2 and SAIV7 VP2. Bottom left: comparisons between GCXV VP4, FNQJV1 VP4 from segments 4-1 and 4-2, FNQJV1 VP4 from segments 4-1 and 4-2, SAIV7 VP4 from segments 2-1 and 2-2. x = segment number, 2, 3 or 4. Last three tables: non statistically significant comparisons.**

| VP4 \ VP1* | GCXV | FNQJV1 x-1 | FNQJV1 x-2 | CTJV1 x-1 | CTJV1 x-2 | SAIV7 x-1 |
|---|---|---|---|---|---|---|
| GCXV | | $8.42 \times 10^{-10}$ | $1.05 \times 10^{-9}$ | $1.65 \times 10^{-9}$ | $1.00 \times 10^{-11}$ | $1.38 \times 10^{-4}$ |
| FNQJV1 x-1 | $7.66 \times 10^{-15}$ | | $5.49 \times 10^{-7}$ | $1.82 \times 10^{-10}$ | $3.34 \times 10^{-10}$ | $3.73 \times 10^{-6}$ |
| FNQJV1 x-2 | $1.59 \times 10^{-12}$ | $8.88 \times 10^{-16}$ | | $2.61 \times 10^{-7}$ | $9.51 \times 10^{-10}$ | $9.58 \times 10^{-4}$ |
| CTJV1 x-1 | $1.89 \times 10^{-15}$ | $1.22 \times 10^{-15}$ | $2.13 \times 10^{-12}$ | | $2.40 \times 10^{-8}$ | $6.80 \times 10^{-4}$ |
| CTJV1 x-2 | $1.72 \times 10^{-11}$ | $6.85 \times 10^{-13}$ | $3.46 \times 10^{-12}$ | 0.00 | | $1.15 \times 10^{-4}$ |
| SAIV7 x-1 | $1.92 \times 10^{-8}$ | $3.25 \times 10^{-8}$ | $8.95 \times 10^{-12}$ | $7.93 \times 10^{-7}$ | $3.23 \times 10^{-7}$ | |
| SAIV7 x-2 | $1.67 \times 10^{-4}$ | $6.86 \times 10^{-4}$ | $1.64 \times 10^{-4}$ | $1.70 \times 10^{-3}$ | $2.77 \times 10^{-4}$ | $6.14 \times 10^{-5}$ |

*For SAIV7, VP2 is the protein used in the alignment with mosquito-derived VP1 sequences (top right).

$p < 0.05$ statistically significant

$p > 0.05$ non statistically significant

| | GCXV VP5-6 | GCXV VP1-3 |
|---|---|---|
| SAIV7 seg2-1 VP4-1 | 0.602 | 0.448 |
| SAIV7 seg2-2 VP4-1 | 0.489 | 0.359 |
| SAIV7 VP2-3 | 0.645 | $1.17 \times 10^{-6}$ |
| GCXV VP1-3 | 0.609 | |

| | GCXV VP6 |
|---|---|
| SAIV7 seg2-1 VP1 | 0.203 |
| SAIV7 seg2-2 VP1 | 0.218 |

| | GCXV VP5 |
|---|---|
| GCXV VP1 | 0.859 |
